# Curing genetic skin disease through altered replication stress response

**DOI:** 10.1101/2020.08.19.255430

**Authors:** Toshinari Miyauchi, Shotaro Suzuki, Masae Takeda, Jin Teng Peh, Masayuki Aiba, Ken Natsuga, Yasuyuki Fujita, Takuya Takeichi, Taiko Sakamoto, Masashi Akiyama, Hiroshi Shimizu, Toshifumi Nomura

## Abstract

Revertant mosaicism, or ‘natural gene therapy’, refers to the spontaneous *in vivo* reversion of an inherited mutation in a somatic cell^1^. Only ∼50 human genetic disorders exhibit revertant mosaicism, implicating a distinctive role played by mutant proteins in somatic correction of a pathogenic germline mutation^2^. However, the process by which mutant proteins induce somatic genetic reversion in these diseases remains unknown. Here we show that heterozygous pathogenic *CARD14* mutations causing autoinflammatory skin diseases, including psoriasis and pityriasis rubra pilaris, are repaired mainly via homologous recombination. Rather than altering the DNA damage response to exogenous stimuli such as X-irradiation or etoposide treatment, mutant CARD14 increased DNA double-strand breaks under conditions of replication stress. Furthermore, mutant CARD14 suppressed new origin firings without promoting crossover events in the replication stress state. Together, these results suggest that mutant CARD14 alters the replication stress response and preferentially drives break-induced replication (BIR), which is generally suppressed in eukaryotes^3^. Our results highlight the involvement of BIR in reversion events, thus revealing a previously undescribed role of BIR that could potentially be exploited to develop therapeutics for currently intractable genetic diseases.

---

Eukaryotic genomes are inevitably challenged by extrinsic and intrinsic stresses that threaten genome integrity. DNA double-strand breaks (DSBs) are the most toxic and mutagenic of DNA lesions^4–6^, arising both exogenously as a consequence of exposure to ionising radiation (IR) and certain chemicals, and endogenously as a result of replication stress or reactive oxygen species. Homologous recombination (HR) is a crucial pathway involved in DSB repair and replication stress response (RSR)^7–9^. HR is generally regarded as a genetically silent event in mitotic cells because it preferentially occurs between sister chromatids following chromosomal replication. However, if HR occurs via a homologous chromosome, it can lead to loss of heterozygosity (LOH), which contributes to cancer initiation and progression^10^, tissue remodelling^11,12^ and, notably, somatic genetic rescue events (referred to as ‘revertant mosaicism [RM]’) in Mendelian diseases^13^. In particular, in some genetic skin diseases, each patient develops dozens to thousands of revertant skin patches where a heterozygous pathogenic germline mutation is reverted via LOH^14–18^, suggesting that HR may be enhanced in mutant keratinocytes, the major cells of the epidermis. Manipulating HR to induce the reversion of disease-causing mutations may potentially benefit patients with various genetic diseases for which only symptomatic treatment is currently available. However, the process by which mutant proteins induce increased rates of HR in these diseases remains unknown.

Caspase recruitment domain-containing protein 14 (CARD14) is a scaffold molecule predominantly expressed in keratinocytes. Heterozygous gain-of-function mutations in *CARD14* lead to CARD14-associated papulosquamous eruption (CAPE), a disease spectrum that includes psoriasis and pityriasis rubra pilaris^19^. Mutant CARD14 (mut-CARD14) triggers the activation of nuclear factor (NF)-κB signalling^20^ with the subsequent disruption of skin homeostasis, inflammation, and hyperproliferation of keratinocytes, which are the canonical pathways implicated in psoriasis^21^.

Here, we present an initial evidence indicating HR-induced RM in CAPE. We aimed to elucidate the molecular mechanisms underlying this self-healing phenomenon by analysing the impact of mut-CARD14 on DNA damage and replication stress, as well as the response to these events. We identified that mut-CARD14 may play an important role in genetic reversion by enhancing BIR under replication stress.

## Results

### Identification of RM in CAPE

We analysed 2 unrelated Japanese patients with CAPE presenting with erythroderma (generalised red skin) since birth. Case 1, a 60-year-old man, and Case 2, a 27-year-old woman^22^, were heterozygous for c.356T>C (p.M119T) and c.407A>T (p.Q136L) in *CARD14*, respectively (Fig. S1a-f). We noted numerous disseminated normal-appearing skin areas in Case 1 and several clinically unaffected spots on the left leg of Case 2 (Fig. 1a and Fig. S1g-k). A skin biopsy revealed that these spots were histologically normalised (Fig. 1b and Fig. S1l-n). Notably, the mutations reverted to wild-type in the epidermis of all examined spots (6 of 6) but not in the dermis (Fig. 1c and Fig. S2a). To address the mechanisms underlying this reversion, we performed a whole-genome oligo-single nucleotide polymorphism (SNP) array analysis using genomic DNA from each epidermis. Notably, copy-neutral LOH (cn-LOH), extending from breakpoints proximal to *CARD14*, to the telomere on chromosome 17q, were identified in 4 of 6 revertant spots (Fig. 1d and Fig. S2b, c). Varying initiation sites of cn-LOH excluded the possibility of simple genetic mosaicism. Notably, 1 of 3 revertant spots in Case 1 showed a different cn-LOH on chromosome 9q (Fig. S2d). Furthermore, Case 1 had developed multiple skin tumours^23,24^, including squamous cell carcinomas (SCCs) which harbour chromosomal aberrations, such as LOH and trisomy (Fig. S2e, f). Because HR events that are generally rare in most genetic skin diseases were frequently observed in these CAPE patients, we inferred that mut-CARD14 may induce an increase in the rate of HR, which potentially leads to RM and carcinogenesis (Fig. S1o).

**Fig. 1.**
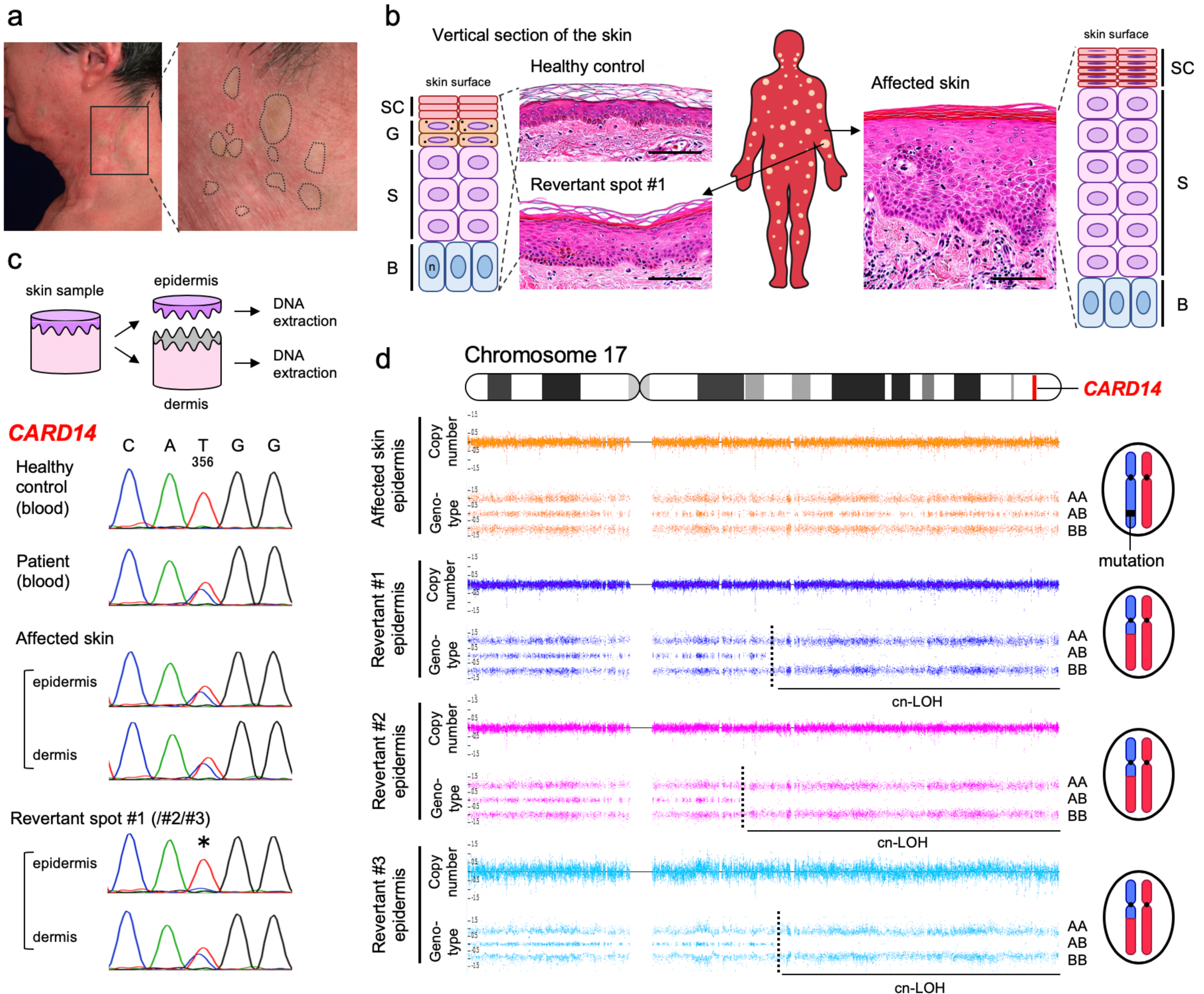
Clinical, histological, and genetic features of Case 1. **a**, Clinically normalised skin spots on the left lateral neck of Case 1. Dotted circles represent the areas suspected of containing revertant spots. **b**, Histological and schematic comparison among healthy control skin, affected skin, and revertant spots. Haematoxylin and eosin staining. Scale bars, 100 µm. SC, stratum corneum; G, granular layer; S, spinous layer; B, basal layer; n, nucleus. See also Fig. S1l-n. **c**, Skin sample processing and mutation analysis of *CARD14* using genomic DNA. The missense mutation, c.356T>C, is absent in the revertant epidermis (*). **d**, SNP array data of chromosome 17 in Case 1. Cn-LOH was identified in all revertant epidermis samples examined in this study. The dotted lines represent recombination breakpoints.

### No direct effect of mutant CARD14 on DDR

To examine the effects of mut-CARD14 on HR, we generated U2OS cell lines which express FLAG-tagged wild-type (wt-), or mut-CARD14, only when doxycycline (Dox) is present (Fig. 2a, b and Fig. S3a-c). To determine whether mut-CARD14 directly increases DSBs, the levels of phosphorylated histone H2AX (γH2AX), a major marker of large chromatin domains surrounding DSBs, were quantified in these cell lines. However, no significant increase in γH2AX levels was detected in any of the cell lines following Dox stimulation (Fig. 2c). To address whether mut-CARD14 alters DNA damage response (DDR) to exogenous DSBs, we monitored γH2AX levels as well as the phosphorylation levels of 53BP1 and RPA2, markers for non-homologous end joining and HR, respectively^25–27^, after exposing the cell lines to IR or etoposide. Mut-CARD14 did not enhance HR in the repair of exogenous DSBs (Fig. S4a-d). To further examine whether mut-CARD14 increases HR in DDR, we performed an enhanced green fluorescent protein (EGFP)-based reporter assay in which a DSB induced by the endonuclease I-SceI followed by HR leads to generation of EGFP-positive cells (Fig. S5a)^28^. Again, neither wt-CARD14 nor mut-CARD14 caused a significant increase in HR frequency (Fig. S5b-f). These findings suggest that mut-CARD14 does not preferentially increase either DNA damage or HR frequency in the DSB repair pathway.

**Fig. 2.**
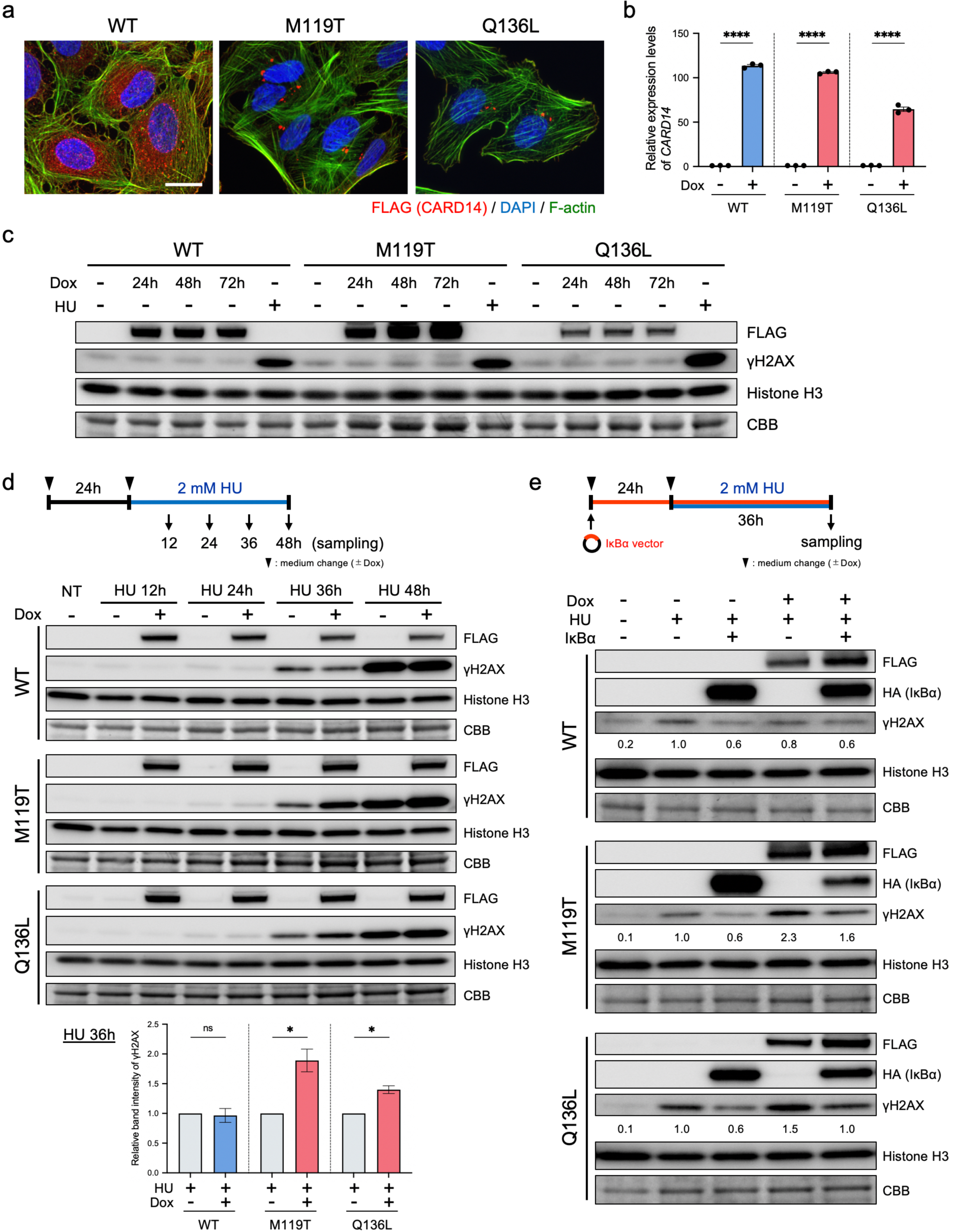
Mut-CARD14 expression alters the RSR. **a**, Intracellular distribution of CARD14 by immunofluorescence. Mut-CARD14 formed aggregates in the perinuclear region, while wt-CARD14 was diffused in the cytoplasm. Scale bar, 25 µm. **b**, Gene expression levels of *CARD14* analysed by qPCR. CARD14 inducible cell lines were incubated in the presence or absence of Dox for 24 h. The results were normalised to *ACTB* expression; n = 3 independent experiments; error bars show s.e.m. **c**, CARD14 inducible cell lines were incubated in the presence of Dox for the indicated time periods. Whole-cell lysate was immunoblotted using the indicated antibodies. HU-treated samples were used as positive controls exhibiting high levels of γH2AX. **d**, Schematic of cells being treated with 2 mM HU, and immunoblots showing the levels of DNA damage following HU treatment. Prolonged HU treatment increased γH2AX in the presence of mut-CARD14. The relative band intensities of γH2AX following 36 h HU treatment are shown in the lower panel; NT, non-treatment. **e**, Whole-cell lysate from cells treated with HU, with or without IκBα overexpression, was immunoblotted with indicated antibodies. IκBα expression partially reduced γH2AX levels in cells expressing mut-CARD14. The band intensities of γH2AX normalised to Histone H3 are also shown. Statistical significance was calculated using two-tailed *t*-test. \**P* < 0.05, \*\*\*\**P* < 0.0001. ns, not significant.

### Mutant CARD14 alters RSR

Replication stress, especially following prolonged stalling, may cause DNA replication fork collapse, leading to DSBs^6^. HR promotes the restart of stress-induced stalled and collapsed replication forks^8^. Therefore, we next sought to address whether mut-CARD14 causes replication stress and alters RSR. To this end, we first performed a fluorescence-activated cell sorting (FACS)-based cell cycle analysis using the U2OS cell lines. However, mut-CARD14 did not alter cell cycle distribution (Fig. S6a, b). This was further confirmed via DNA fibre analysis, in which mut-CARD14 neither affected fork speed nor increased stalled fork frequency (Fig. S6c-g). Thus, mut-CARD14 did not result in replication stress per se.

To address whether mut-CARD14 increases DSBs under conditions of replication stress, we treated the cell lines with hydroxyurea (HU) and monitored γH2AX levels up to 48 h after treatment. Notably, mut-CARD14 cell lines with Dox showed higher levels of γH2AX than those without Dox at 36 h and 48 h, whereas wt-CARD14 cell lines showed no difference at all time points (Fig. 2d). We performed the same analysis using aphidicolin (APH), which also caused similar differences in γH2AX levels based on mut-CARD14 expression (Fig. S6h). In both analyses, no obvious difference in γH2AX levels was detected 24 h following treatment with HU or APH. Collectively, these results suggest that mut-CARD14 expression leads to fork instability following prolonged replication fork stalling but does not increase DSBs following short-term fork stalling.

We next sought to clarify whether mut-CARD14-induced stalled fork instability is mediated by the NF-κB pathway, which is activated by mut-CARD14^20^ (Fig. S6i). Therefore, we examined γH2AX levels in HU-treated cell lines with or without an inhibitor of κB (IκB) overexpression which suppresses NF-κB activation (Fig. S6j). Notably, IκBα overexpression partially reduced γH2AX levels in the cells expressing mut-CARD14 (Fig. 2e). Mut-CARD14-induced alteration of the RSR is partially explained by activation of the NF-κB signalling pathway.

### HR activation by mutant CARD14

Having confirmed that mut-CARD14 expression decreases fork stability under prolonged replication stress, we investigated whether HR-related factors are activated under those circumstances. We approached this issue using a FACS-based method to quantify the activities of HR-related factors at the single cell level^29,30^. For these analyses, the mean values of signal intensities obtained from HU-treated and Dox-absent samples were set to threshold levels and the percentages of cells showing intensities above these levels (γH2AX^hi^ cells) were compared among samples (Fig. 3a, b and Fig. S7a). Notably, consistent with immunoblotting data, mut-CARD14 cell lines treated with Dox and either HU or APH for 36 h, showed a significant increase in the percentage of γH2AX^hi^ cells to the total cells, compared to those without Dox, whereas wt-CARD14 showed no increase (Fig. 3c and Fig. S7b-d). Similarly, under prolonged replication stress, mut-CARD14 expression significantly increased the ratio of highly phosphorylated RPA2 cells (p-RPA2^hi^ cells) (Fig. 3d). Furthermore, co-staining with γH2AX and p-RPA2 revealed that mut-CARD14 expression increased the proportion of double-positive cells (γH2AX^hi^p-RPA2^hi^ cells) (Fig. 3b, e). Similarly, we identified that mut-CARD14 also activated BRCA1, which is important for HR and fork restart^31^, yielding a higher proportion of γH2AX^hi^p-BRCA1^hi^ cells (Fig. 3f, g). We also examined whether the ATR-CHK1 pathway, which stabilises replication forks and promotes several types of fork restarts including a HR-mediated fork restart in RSR^32^, is activated by mut-CARD14 under prolonged fork stalling. Mut-CARD14 significantly increased the proportion of γH2AX^hi^p-CHK1^hi^ cells as well as p-CHK1^hi^ cells (Fig. 3h, i). Collectively, these results suggest that in mut-CARD14-expressing cells, collapsed forks caused by prolonged stalling were repaired by the HR-mediated pathway via activation of the ATR-CHK1 signalling pathway.

**Fig. 3.**
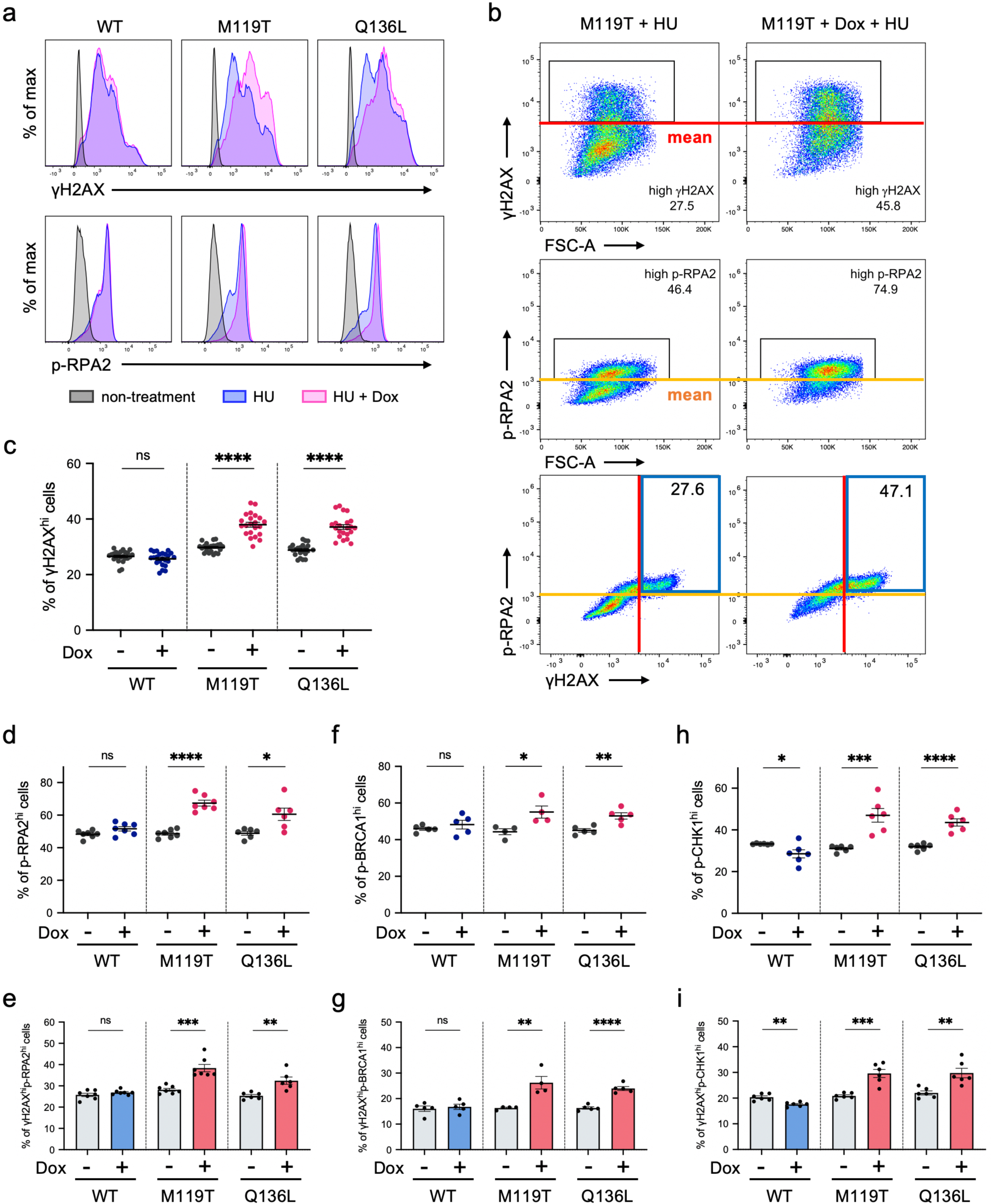
FACS-based analyses of HR-related factors. **a**, Expression of γH2AX (upper panels) and p-RPA2 (lower panels) in CARD14 inducible cell lines. Each panel is a representative histogram showing comparison between non-treatment and 2 mM HU-treatment for 36 h with or without Dox. **b**, Red and yellow lines are the threshold levels which were determined based on the mean values of signal intensities obtained from HU-treated and Dox-absent cells. The numbers in the upper and middle panels represent the percentage of γH2AX^hi^ or p-RPA2^hi^ cells, and those in the blue boxes in the lower panels represent γH2AX^hi^p-RPA2^hi^ cells. **c**, Comparison of the percentage of γH2AX^hi^ cells; n = 23 (WT) or 22 (M119T/Q136L) independent experiments. Error bars show s.e.m. **d, f, h**, Comparison of the percentage of p-RPA2^hi^ (**d**), p-BRCA1^hi^ (**f**), and p-CHK1^hi^ cells (**h**). **e, g, i**, Comparison of the percentage of γH2AX^hi^p-RPA2^hi^ cells (**e**), γH2AX^hi^p-BRCA1^hi^ cells (**g**), and γH2AX^hi^p-CHK1^hi^ cells (**i**). n = 4-7 independent experiments. Error bars represent s.e.m. Statistical significance was calculated using the two-tailed t-test. \**P* < 0.05, \*\**P* < 0.01, \*\*\**P* < 0.001, \*\*\*\**P* < 0.0001; ns, not significant.

### Mutant CARD14 promotes BIR in RSR

HR-mediated repair of stalled or collapsed forks may take place via 3 pathways: synthesis-dependent strand annealing (SDSA), double Holliday junction (dHJ), and BIR^3,9^ (Fig. S8a). BIR is initiated when only one broken end is available for strand invasion, whereas SDSA and dHJ require two DSB ends. Thus, dormant origin firing is essential for rescuing collapsed forks using these two pathways^3^. We sought to determine which pathway played a dominant role in replication fork repair under mut-CARD14 expression. To this end, we first performed quantitative PCR (qPCR) analysis of replication-related genes. The expression levels of genes associated with replication origin firing, such as *ORC1, MCM10, GINS1*, and *GINS2*, were decreased in mut-CARD14-expressing cells following extended HU treatment, but not in wt-CARD14-expressing cells (Fig. S9a-h). To further confirm whether mut-CARD14 suppresses new origin firing after prolonged fork stalling, we analysed replication fork dynamics via DNA fibre assay with prolonged HU treatment. The frequency of new origin firing was reduced by mut-CARD14 expression but not by wt-CARD14, while neither wt-nor mut-CARD14 altered the frequency of stalled or ongoing replication forks (Fig. 4a-d). As these findings suggested that mut-CARD14 expression, together with HU, inhibits dormant origin firings near collapsed forks, resulting in a situation where only one broken end is available for DSB repair, we concluded that mut-CARD14 promotes BIR in stalled or collapsed fork repair under conditions of replication stress (Fig. S8b).

**Fig. 4.**
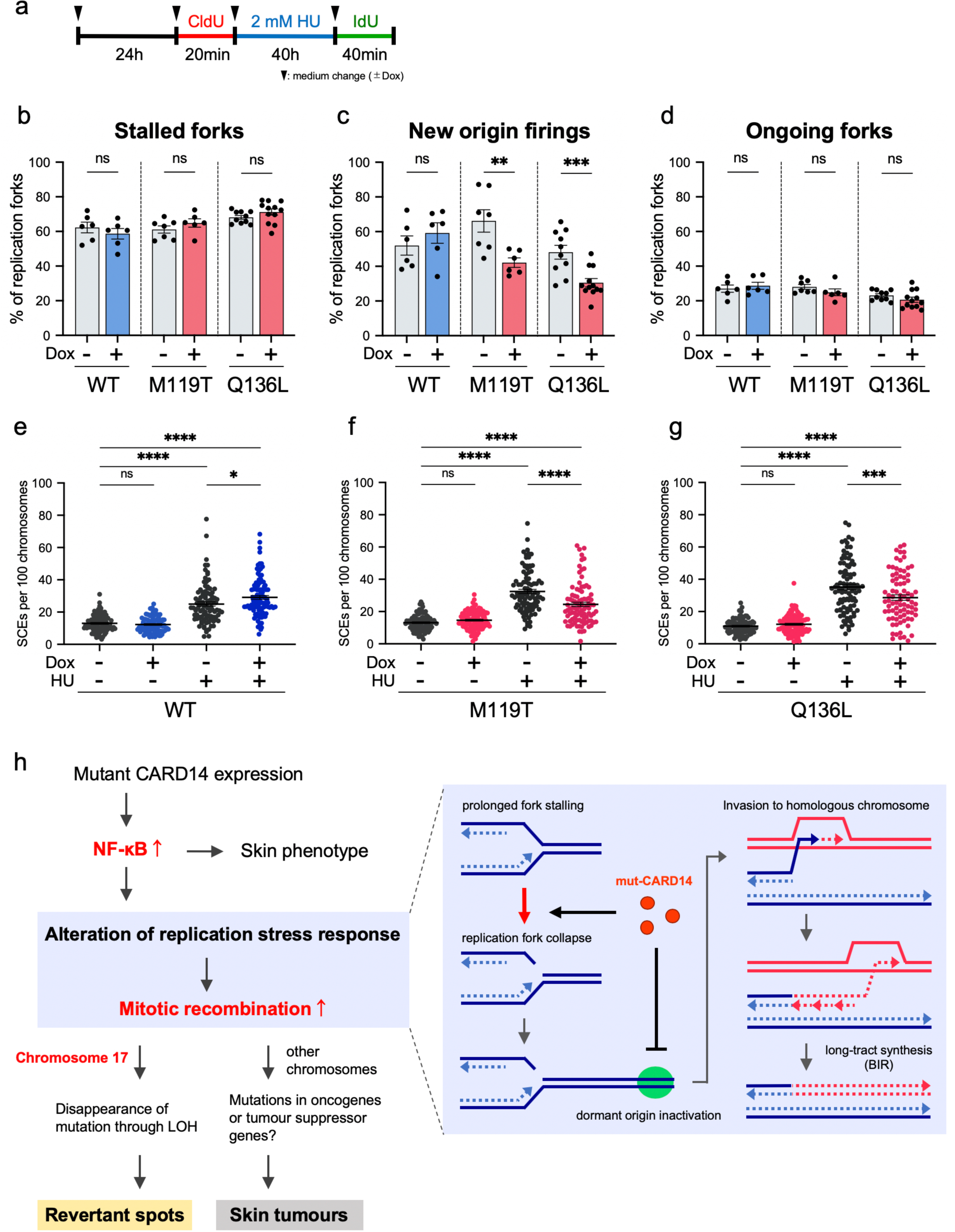
Mut-CARD14 expression suppresses dormant origin firings and promotes BIR. **a**, Schematic depicting timeline of when cells were labelled with CldU and IdU in the presence of HU treatment. **b, c, d**, Quantification of the percentage of stalled forks (**b**), new origin firings (**c**), and ongoing forks (**d**) to all CldU-labelled forks; n = 6-12 independent experiments; error bars show s.e.m. Statistical significance was calculated using the two-tailed t-test. **e, f, g**, Quantification of SCE formation in cells treated with or without HU. Quantification of SCE formation in cells treated with or without 2 mM HU for 36 h. For each condition, 27-35 metaphase cells were analysed and each data point represents the number of SCEs per 100 chromosomes per metaphase spread. Data from 3 experiments were pooled. Horizontal lines represent mean values ± s.e.m. Statistical significance was calculated using one-way ANOVA followed by a multiple comparisons test. **h**, A model of the mechanism underlying mitotic recombination induced by mut-CARD14 expression. \**P* < 0.05, \*\**P* < 0.01, \*\*\**P* < 0.001, \*\*\*\**P* < 0.0001; ns, not significant.

Notably, HR-mediated revertant epidermis samples as well as SCC samples, obtained from the studied CAPE patients, all possessed long-tract LOH extending from recombination sites to the telomere (Fig. 1d and Fig. S2b, d-f). Although this specific form of LOH arises only via BIR or dHJ with crossover resolution (Fig. S8c), other possible consequences of dHJ, such as gene conversion, were not detected in any of the samples. Furthermore, the frequency of sister chromatid exchanges (SCEs)^33^, which represents the crossover events, following HU treatment was somewhat decreased in mut-CARD14-expressing cells, but not in those expressing wt-CARD14 (Fig. 4e-g), suggesting that dHJ with crossover does not mainly occur in this context. Considering the results together, we deduced that mut-CARD14 plays a role in the reversion of a disease-causing mutation in patients with CAPE, by enhancing BIR under conditions of replication stress (Fig. 4h).

## Discussion

The current study revealed, for the first time, that RM occurs in CAPE. We demonstrated that HR is the major mechanism responsible for the reversion of *CARD14* mutations and that mut-CARD14 preferentially drives BIR under conditions of replication stress. However, BIR is generally regarded as a rare event which is suppressed by MUS81 endonuclease in normal cells, as well as converging forks arriving from the opposite direction, as it elevates the risk of mutagenesis and chromosomal abnormalities^34^. Notably, recent studies have uncovered a series of unusual BIR-promoting circumstances as follows^3,35–40^: (i) generation of DNA nicks in genomic regions lacking replication origins within a MUS81-null background, (ii) repair of massive replication fork collapse under conditions of replication stress (e.g., overexpression of oncogenes, deregulation of origin licensing, inverted DNA repeats, trinucleotide repeats, and R-loops), and (iii) alternative lengthening of telomeres in the absence of telomerase. Our findings suggest that mut-CARD14 promotes replication fork collapse and suppresses dormant origin firing in the replication stress state. These effects of mut-CARD14 collectively promote BIR that may continue for hundreds of kilobases, resulting in long-tract LOH that extends to the telomere. Therefore, to our knowledge, this study is the first to provide evidence indicating the potential contribution of BIR to RM. Notably, similar long-tract LOH has also been frequently observed in other skin diseases such as ichthyosis with confetti and loricrin keratoderma where dozens to thousands of HR-driven revertant skin spots arise in each patient^14-18^. Further investigations centred on RM may help elucidate hitherto unknown or unvalued mechanisms that regulate the BIR pathway.

This study also suggests that skin inflammation such as that induced by mut-CARD14 may promote BIR. Recent studies on patients with ulcerative colitis have revealed that human intestinal stem cells exposed to long-standing inflammation adapt to such inflammation by acquiring genetic and genomic alterations including LOH, associated with the downregulation of IL-17 signalling^11,12^. Furthermore, long-tract LOH is frequently seen in skin lesions of porokeratosis, a common autoinflammatory keratinisation disease^41^. Interestingly, previous studies have shown that it is replication stress, and not I-SceI-induced DSB, which activates the NF-κB signalling pathway, which, in turn, induces HR^42^. The current study indicated that mut-CARD14 does not promote HR when DSBs are exogenously introduced via I-SceI but drives BIR under conditions of replication stress. These findings suggest that inflammation resulting from NF-κB activation under conditions of replication stress may induce BIR. Further studies are warranted to address the involvement of inflammation in BIR initiation.

In conclusion, this study demonstrated that RM occurs in CAPE and implicates the involvement of BIR in reversion events. Fully elucidating the molecular mechanisms underlying natural gene therapy may deepen the understanding of DDR or RSR and pave the way to develop innovative therapies for genetic diseases, for which therapeutic options are currently limited, by manipulating HR to repair disease-causing mutations.

## Methods

### Human subjects and study approval

Patients participating in the study provided written informed consent, in compliance with the Declaration of Helsinki. This study was approved by the Institutional Review Board of the Hokkaido University Graduate School of Medicine (project No. 14-063). Both patients provided a peripheral blood or saliva sample and skin samples from affected skin and revertant spots.

### Genomic DNA extraction

For mutation analysis, genomic DNA from the patients’ peripheral blood or saliva was extracted using a QIAamp DNA Blood Maxi Kit (Qiagen) or an Oragene DNA Self-Collection Kit (DNA Genotek), respectively. Genomic DNA was also extracted from the patients’ skin using a QIAamp DNA Micro Kit (Qiagen) after separating the epidermis from the dermis of punch-biopsied skin samples using an ammonium thiocyanate solution, as described previously^43^.

### Sanger sequencing

Exons and exon-intron boundaries in *CARD14* were amplified via PCR using AmpliTaq Gold PCR Master Mix (ThermoFisher Scientific). Primer sequences and PCR conditions are available upon request. PCR amplicons were treated with ExoSAP-IT reagent (Affymetrix), and the sequencing reaction was performed using BigDye Terminator version 3.1 (ThermoFisher Scientific). Sequence data were obtained using an ABI 3130xl genetic analyser (Applied Biosystems).

### Whole-genome oligo-SNP array

CytoScan HD Array (Affymetrix) was used to identify copy number variations (CNVs) and LOHs using genomic DNA extracted from the epidermis (outsourced to RIKEN Genesis). All experimental procedures were conducted according to the manufacturer’s instructions. Briefly, 250 ng of each genomic DNA sample was digested with Nsp I and ligated to adaptors, followed by PCR amplification. Following purification, fragmentation, and biotinylation of the PCR products, these samples were hybridised to the Affymetrix CytoScan HD Array. After washing and staining, fluorescent signals were obtained with a GeneChip Scanner 3000 7G (Affymetrix) and GeneChip Command Console (Affymetrix). The data were then analysed using Chromosome Analysis Suite Software 4.0 (Affymetrix) by filtering CNVs with a minimum size of 400 kbp and 50 consecutive markers and LOHs with a minimum size of 3,000 kbp and 50 consecutive markers.

### Cell culture, X-irradiation, and drug treatment

U2OS (ATCC), the human osteosarcoma cell line, and HaCaT (CLS Cell Lines Service), the commercially available immortalised human keratinocyte cell line, were cultured in Dulbecco’s modified Eagle medium (Nacalai Tesque) supplemented with 10% foetal bovine serum (Sigma-Aldrich) and with or without 1% Antibiotic Antimycotic Solution (Sigma-Aldrich). To examine the response to X-irradiation, 150 kV X-ray was delivered at 20 mA with 1 mm aluminium filter at a distance of 350 mm from the cell surface. To analyse the response to etoposide, cells were treated with 10 or 250 µM etoposide (Sigma-Aldrich) for indicated time periods since etoposide was reported to induce two types of DNA damage mechanisms depending on its concentration^44^. To investigate the response to HU and APH, cells were treated with 2 mM HU (Sigma-Aldrich) and 2 µg/mL APH (Sigma-Aldrich) for indicated time periods, respectively. The cells were then washed with phosphate-buffered saline (PBS) (Nacalai Tesque) before addition of drug-free medium and incubated as necessary.

### RNA extraction and quantitative real-time reverse transcription PCR

RNA extraction and reverse transcription were performed using a SuperPrep II Cell Lysis & RT Kit for qPCR (Toyobo). Quantitative real-time PCR was carried out using the QuantStudio 12K Flex Real Time PCR System (ThermoFisher Scientific) with TaqMan Fast Advanced Master Mix (ThermoFisher Scientific) and TaqMan MGB probes (ThermoFisher Scientific) according to the manufacturer’s instructions. To assess the expression levels of CARD14 and inflammatory chemokines, the following TaqMan probes were used: *CARD14* (Hs01106900_m1), *CXCL8* (Hs00174103_m1), and *CCL20* (Hs00355476_m1). For the analysis of replication-related genes, Custom TaqMan Array Cards (ThermoFisher Scientific) were used. Target genes and their corresponding TaqMan probes were as follows; *ORC1* (Hs01069758_m1), *ORC6* (Hs00941233_g1), *MCM2* (Hs01091564_m1), *MCM7* (Hs00428518_m1), *MCM10* (Hs00218560_m1), *CDC6* (Hs00154374_m1), *CDC45* (Hs00907337_m1), *CDT1* (Hs00925491_g1), *TOP3A* (Hs00172806_m1), *TOP3B* (Hs00172728_m1), *TOPBP1* (Hs00199775_m1), *RECQL4* (Hs00171627_m1), *GINS1* (Hs01040834_m1), *GINS2* (Hs00211479_m1), *GINS3* (Hs01090589_m1), *GINS4* (Hs01077879_m1), *TIPIN* (Hs00762756_s1), *CLSPN* (Hs00898637_m1), *WDHD1* (Hs00173172_m1), *PCNA* (Hs00427214_g1), *POLA1* (Hs00213524_m1), *POLD1* (Hs01100821_m1), *POLE* (Hs00923952_m1), *BOD1L1* (Hs00386033_m1), and *RFC1* (Hs01099126_m1). Following comprehensive analyses, expression levels of some genes were confirmed using separately prepared samples. Expression values were normalised to *ACTB* (Hs01060665_g1) levels, and relative expression levels were calculated using the ΔΔCt method.

### Plasmids and transfection

Wild-type *CARD14* cDNA (pFN21AE3344) was purchased from the Kazusa DNA Research Institute. To generate CARD14-expressing vector (p3FLAG-CARD14wt) for transient overexpression, *EGFP* in pEGFP-C2 (Takara Bio) was replaced by the wild-type *CARD14* gene obtained from pFN21AE3344 with N-terminal 3xFLAG tag peptides (DYKDHDGDYKDHDIDYKDDDDK) (Sigma-Aldrich). To generate CARD14-expressing vector (pTRE3G-Pur-3FLAG-CARD14wt) for establishment of Tet-On 3G inducible cell lines, the wild-type CARD14 insert with N-terminal 3xFLAG tag was cloned into pBApo-EFalpha Pur DNA (Takara Bio) whose EF-1α promoter was replaced by TRE3G promoter obtained from pTRE3G (Clontech). The c.356T>C or c.407A>T mutation was introduced both into p3FLAG-CARD14wt and pTRE3G-Pur-3FLAG-CARD14wt using QuikChange Lightning Site-Directed Mutagenesis Kits (Agilent Technologies) (p3FLAG-CARD14-M119T /-Q136L or pTRE3G-Pur-3FLAG-CARD14-M119T /-Q136L, respectively). To establish Tet3G-expressing U2OS cells, pCMV-Tet3G (Clontech) was used. For the NF-κB-Luciferase Reporter assay, pNL1.1.PGK[*Nluc*/PGK] (Promega), pGL4.27[*luc2P*/minP/Hygro] (Promega), and pNL3.2.NF-κB-RE[*NlucP*/NF-κB-RE/Hygro] (Promega) were purchased, and pCMV4-3 HA/IkB-alpha was a gift from Warner Greene (Addgene, #21985). To generate firefly luciferase (Fluc)-expressing vector (pNL1.1.PGK[*Fluc*/PGK]) for transfection efficiency normalisation, *Nanoluc luciferase* (Nluc) gene in pNL1.1.PGK[*Nluc*/PGK] was replaced by *Fluc* gene obtained from pGL4.27[*luc2P*/minP/Hygro]. pDRGFP (Addgene, #26475) and pCBASceI (Addgene, #26477) were gifts from Maria Jasin for DR-GFP assay. To generate wild-type EGFP-expressing vector (pDRGFPwt) for transfection efficiency normalisation, *Sce-EGFP* gene, which contains an I-SceI recognition sequence, was replaced by *EGFP* gene obtained from pEGFP-C2. All plasmids were transfected into U2OS or HaCaT cells using Lipofectamine 3000 (ThermoFisher Scientific) or Xfect Transfection Reagent (Takara) according to the manufacturer’s instructions.

### Establishment of the Tet-On 3G CARD14 inducible cell lines

The Tet-On 3G Inducible Expression System (Clontech) was used according to the manufacturer’s instructions to establish Tet-On 3G CARD14 inducible cell lines. Briefly, U2OS cells were first transfected with the pCMV-Tet3G plasmid and selected using 250 µg/ml G418 (Sigma-Aldrich). The G418-resistant clone was maintained as the Tet3G-expressing U2OS cell line. Tet3G-expressing U2OS cells were then transfected with pTRE3G-Pur-3FLAG-CARD14wt, pTRE3G-Pur-3FLAG-CARD14-M119T, or pTRE3G-Pur-3FLAG-CARD14-Q136L. Next, 0.75 µg/µl puromycin (Gibco) was used for selection following which double-stable Tet-On 3G inducible cell lines were cloned by limiting the dilution technique. The cell lines of each genotype were tested for CARD14 expression in the presence of 500 ng/ml Dox (Clontech), and clones with the highest fold induction were selected for further analyses.

### Immunofluorescence staining and microscopy

Twenty-four hours after adding Dox into CARD14 inducible cell lines, or 24 h after transfection into U2OS or HaCaT cells, cells were fixed in 4% paraformaldehyde in PBS for 10 min at 4°C and washed twice with PBS. After permeabilization with 0.5% Triton X-100 in PBS for 10 min at 4°C and blocking with 3% bovine serum albumin (BSA) (Wako) in PBS with 1% DAPI (Sigma-Aldrich) at 37°C, the cells were incubated with primary and secondary antibodies diluted with 3% BSA in PBS for 60 min (primary antibodies) or for 30 min (secondary antibodies) at 37°C. Anti-FLAG M2 antibodies (Sigma-Aldrich, F3165) were used as the primary antibodies for FLAG staining. Alexa Fluor 488-conjugated goat anti-mouse IgG (H + L) (ThermoFisher Scientific, A11001) and Alexa Fluor 647-conjugated goat anti-mouse IgG (H + L) (ThermoFisher Scientific, A21244) were used as secondary antibodies. For F-actin staining, the cells were incubated with Alexa Fluor 488 Phalloidin (ThermoFisher Scientific) for 20 min at 37°C between the primary and secondary antibody reactions. Nuclei were stained with DAPI during the blocking step. Fluorescence images were obtained with a BZ-X800 (Keyence) or FV1000 confocal laser scanning microscope (Olympus).

### NF-κB-Luciferase Reporter assay

CARD14 inducible cell lines were co-transfected with pNL3.2.NF-κB-RE[*NlucP*/NF-κB-RE/Hygro] and pNL1.1.PGK[*Fluc*/PGK] in the absence or presence of Dox. To evaluate the effect of IκBα expression on the NF-kB signalling pathway, pCMV4-3 HA/IkB-alpha was overexpressed using pNL3.2.NF-κB-RE[*NlucP*/NF-κB-RE/Hygro] and pNL1.1.PGK[*Fluc*/PGK]. Luciferase activity of the lysates was measured using a Nano-Glo Dual-Luciferase Reporter Assay System (Promega) and SpectraMax Paradigm (Molecular Devices) according to the manufacturer’s instructions. Nluc activity derived from pNL3.2.NF-κB-RE[*NlucP*/NF-κB-RE/Hygro] was normalised to Fluc activity from pNL1.1.PGK[*Fluc*/PGK].

### Western blotting

The whole cell lysate was obtained by lysing cells in a mixture of NuPAGE LDS Sample Buffer (ThermoFisher Scientific) and NuPAGE Sample Reducing Agent (ThermoFisher Scientific). The lysate was sonicated with Handy Sonic UR-21P (TOMY), denatured for 10 min at 70°C, and fractionated on NuPAGE 4-12% Bis-Tris Protein Gels (ThermoFisher Scientific), followed by the transferring of resolved proteins onto PVDF membranes via an iBlot 2 Dry Blotting System (ThermoFisher Scientific). Following blocking for 60 min with 1% BSA in TBST (10 mM Tris-HCl (pH7.4), 150 mM NaCl, 0.05% Tween 20) or 5% skim milk in TBST, membranes were incubated overnight at 4°C with one of the following primary antibodies: FLAG (Sigma-Aldrich, F3165), γH2AX (Millipore, 05-636), histone H3 (abcam, ab1791), RPA32 (RPA2) (Santa Cruz Biotechnology, sc-56770), phospho-RPA2 (Ser33) (Novus Biologicals, NB100-544), 53BP1 (Cell Signaling Technology, 4937), phospho-53BP1 (Ser1778) (Cell Signaling Technology, 2675), BRCA1 (Santa Cruz Biotechnology, sc-6954), or phospho-BRCA1 (Ser1524) (Cell Signaling Technology, 9009). HRP-linked horse anti-mouse IgG (Cell Signaling Technology, 7076) and HRP-linked goat anti-rabbit IgG (Cell Signaling Technology, 7074) were used as secondary antibodies. 20X LumiGLO Reagent and 20X Peroxide (Cell Signaling Technology) or SuperSignal West Dura Extended Duration Substrate (ThermoFisher Scientific) were used as a substrate for the secondary antibodies. Chemiluminescence data were obtained with ImageQuant LAS 4000 (Fujifilm) and band intensities were quantified using Image J software (NIH).

### DR-GFP assay (I-SceI induced DSB repair assay)

CARD14 inducible cell lines were used in a DR-GFP assay via transient transfection with pDRGFP and pCBASceI. Cells were harvested 72 h after transfection with these two plasmids, following which the number of EGFP-positive cells was measured using FACSCanto II (BD Biosciences) (30,000 cells per biological replicate were analysed). Under these conditions, parallel transfection with the pDRGFPwt plasmid was used to determine transfection efficiency. All analyses were performed in the absence or presence of Dox. Cells not transfected with pCBASceI were used as negative control. A DR-GFP assay using U2OS cells which harbour chromosomally integrated EGFP reporter substrates was also conducted. To establish this cell line, U2OS cells were first transfected with pDRGFP plasmid, selected using 0.75 µg/µl puromycin and cloned via the limiting dilution technique (U2OS_DRGFP cells). U2OS_DRGFP cells were transfected with p3FLAG-CARD14-WT, p3FLAG-CARD14-M119T, or p3FLAG-CARD14-Q136L along with pCBASceI. Cells were harvested 72 h after transfection and the number of EGFP-positive cells were counted using FACSCanto II. HR efficiency was calculated as the ratio of EGFP-positive cells over total cells. Non-transfected cells were used as negative control. All flow cytometry data were analysed using FlowJo software (BD Biosciences).

### Cell cycle analysis by FACS

To analyse cell cycle distribution, cells were pulse labelled with 10 µM EdU 30 min before harvest. EdU staining was performed using a Click-iT Plus EdU Alexa Fluor 647 Flow Cytometry Assay Kit (ThermoFisher Scientific) according to the manufacturer’s instructions. Genomic DNA was stained with FxCycle Violet (ThermoFisher Scientific). Flow cytometry data of EdU-DNA content profiles were acquired with FACSCanto II and analysed using FlowJo software.

### FACS-based quantification of DNA damage and HR-related factor activity

Cells treated with or without HU were trypsinised and washed with 1% BSA in PBS and fixed in 4% paraformaldehyde in PBS for 15 min at room temperature, followed by incubation with dilution buffer (0.5% BSA, 0.1% Triton X-100 in PBS) for 30 min at room temperature. Cells were then incubated for 30 min at room temperature with one or two of the following primary antibodies diluted with dilution buffer: γH2AX (Millipore, 05-636), phospho-RPA2 (Ser33) (Novus Biologicals, NB100-544), phospho-BRCA1 (Ser1524) (Cell Signaling Technology, 9009), or phosphor-Chk1 (Ser345) (abcam, ab47318). After washing with dilution buffer, cells were subsequently incubated with Alexa Fluor 488-conjugated goat anti-mouse IgG (H + L) and/or Alexa Fluor 647-conjugated goat anti-rabbit IgG (H + L) as secondary antibodies. Genomic DNA was stained with FxCycle Violet. A gate was established for each factor using negative control samples stained with mouse IgG1 negative control antibodies (BIO-RAD, MCA928) or polyclonal rabbit IgG (R&D Systems, AB-105-C). In all analyses, signal intensities of 30,000 cells per biological replicate were measured using FACSCanto II and analysed with FlowJo software.

### DNA fibre assay

The DNA fibre assay was performed as described previously^45^ with slight modifications. Briefly, CARD14 inducible cell lines were labelled with 25 µM 5-chlorodeoxyuridine (CldU) (Sigma-Aldrich), washed with PBS, and exposed to 250 µM 5-iododeoxyuridine (IdU) (Sigma-Aldrich). To measure the efficiency of replication restart, cells were treated with 2 mM HU after CldU labelling and exposed to IdU. Labelled cells were harvested and resuspended in cold PBS. The cell suspension was mixed 1:4 with cell lysis solution (0.5% SDS, 50 mM EDTA, 200 mM Tris-HCl pH 8.0), placed on glass slides and carefully tilted at a 15° angle causing DNA fibres to spread into single molecules via gravity. Next, the DNA fibres were denatured by fixing with 4% paraformaldehyde in PBS and immersing in 2.5 M HCl for 80 min at 27°C. Slides were neutralised and washed with PBS before blocking with 3% BSA in PBS for 30 min at 37°C. DNA fibres were then incubated with primary and secondary antibodies diluted in 3% BSA in PBS overnight at 4°C (primary antibodies) or for 90 min at 37°C (secondary antibodies). Anti-BrdU antibodies [BU1/75 (ICR1)] (abcam, ab6326) for CldU and anti-BrdU antibodies [clone B44] (BD Biosciences, 347580) for IdU were used as primary antibodies. Alexa Fluor 488-conjugated goat anti-rat IgG (H + L) antibodies and Alexa Fluor 594-conjugated goat anti-mouse IgG (H + L) antibodies (ThermoFisher Scientific, A11005) were used as secondary antibodies. Finally, slides were mounted in ProLong Diamond Antifade Mountant (ThermoFisher Scientific). Images were obtained using a FV1000 confocal laser scanning microscope (Olympus) and all data were analysed via Computer Assisted Scoring & Analysis (CASA) software purchased from Dr. Paul Chastain. Tracts containing CldU were pseudocoloured in red and tracts containing IdU were green. Replication fork speed was estimated using a conversion factor of 2.59 kb/mm^46^.

### SCE assay

The SCE assay was carried out essentially as previously described^47^. Prior to the experiment, slides were immersed in 0.1 N HCl in 99.5% ethanol for 20 min at room temperature and washed thrice with 99.5% ethanol. After rinsing with distilled water, they were stored in distilled water at 4°C. Cells were incubated with 20 µM BrdU (Sigma-Aldrich) for 42 h. During the last 2 h of BrdU incubation, cells were also treated with 0.2 µg/ml colcemid (Sigma-Aldrich). To examine the effects of replication stress, cells were incubated with BrdU for 20 h followed by 2 mM HU treatment for 36 h, incubated again with BrdU for 24 h and treated with colcemid for the last 2 h. Cells were collected by mitotic shake-off, washed in PBS and swollen in 7 ml of hypotonic solution (46.5 mM KCl, 8.5 mM Na⋅Citrate) for 13 min at 37 °C. After the incubation, 2 ml of freshly prepared 3:1 methanol-acetic acid fixative was added and mixed by gently inverting the tube. The tubes were centrifuged for 3 min at 200 x g at room temperature and the supernatant was aspirated. The cell pellets were resuspended with 4 ml of fixative and incubated for 20 min at 4°C. Following fixation, cells were resuspended in a minimal amount of fixative. Metaphase cells were then spread on the chilled slides described above and completely dried at room temperature. Slides were stained for 30 min at room temperature with 10 µg/ml Hoechst 33258 (Sigma-Aldrich) in Sorensen’s phosphate buffer (0.1 M Na_2_HPO_4_, 0.1 M KH_2_PO_4_, pH 6.8). After washing with Sorensen buffer, the slides were covered with Sorensen buffer and exposed to UV light for 60 min at 55°C, following which the slides were immersed in 1x SSC buffer (ThermoFisher Scientific), incubated for 60 min at 50°C and finally stained with 10% Gimsa (Wako) in Sorensen buffer for 30 min at room temperature. After washing thrice with water, the coverslips were mounted on the slides using MOUNT-QUICK (Funakoshi). Images were obtained via BZ-X800 (Keyence) using CFI Plan Apo λ 100xH objective lens. At least 25 images of each condition were randomly taken and the number of SCEs per chromosome was scored.

### Quantification and statistical analysis

Statistical parameters are shown in the figures and legends. Two-tailed t-tests or one-way ANOVA with multiple comparisons tests were used for comparisons of means of normally distributed data, whereas two-tailed Mann-Whitney U tests were used for comparison of non-normally distributed data. All statistical analyses were conducted using Prism 8.0 (GraphPad) and the threshold for defining statistical significance was *P* < 0.05.

## Acknowledgements

We are most indebted to the patients and their family members for their participation in this study. This work was supported by the JSPS KAKENHI (Grant numbers: JP19H03679 and JP17H06271 to T.N. and H.S., respectively), the Terumo Foundation for Life Sciences and Arts (Grant number: 16-II 330 to T.N.), the Rohto Dermatology Research Award (to T.N.), the Akiyama Life Science Foundation (to T.N.), the Nakatomi Foundation (to T.N.), the Ichiro Kanehara Foundation (to T.N.), the Takeda Science Foundation (to T.N.), the Geriatric Dermatology Research Grant (to T.N.), and the Northern Advancement Center for Science & Technology (NOASTEC) Foundation (Grant number: H28 T-1-42 to T.N.).

## Author Contributions

T.M, S.S., and T.N. designed the study. K.N., Y.F., T.T., T.S., M. Ak., H.S., and T.N. provided specimens. T.M. conducted the majority of the experiments and data analyses. S.S. assisted with mutation analysis. S.S. and M.T. assisted with the DNA fibre assay. M.T., J.T.P, and M.Ai. assisted with Western blot analysis. T.M. and T.N. wrote the manuscript.

