## Supplemental data for "Curing genetic skin disease through altered replication stress response"

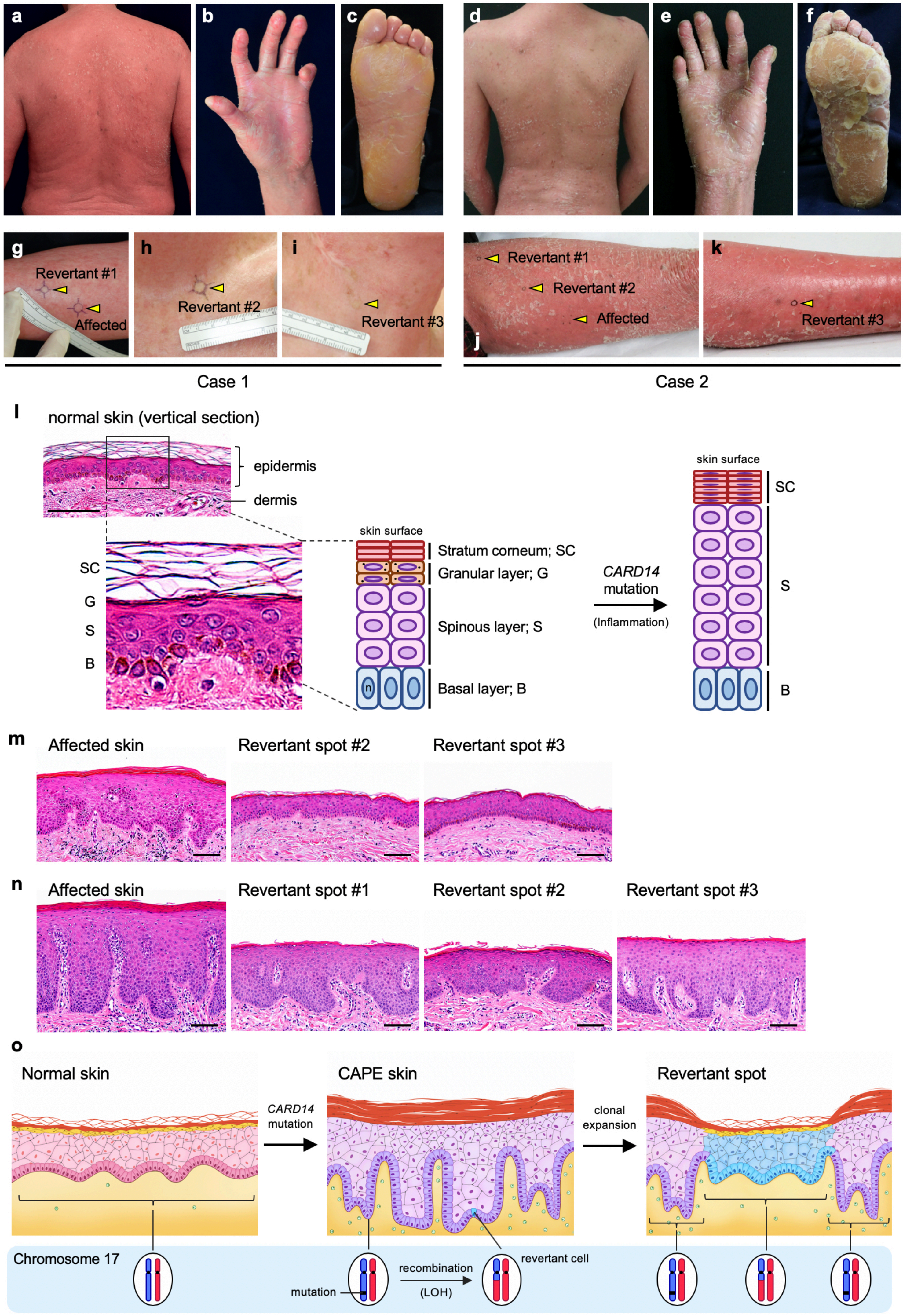

a
b
c
d
e
f
g
Revertant #1
Affected
h
Revertant #2
i
Revertant #3
j
Revertant #1
Revertant #2
Affected
k
Revertant #3
Case 1
Case 2
l
normal skin (vertical section)
epidermis
dermis
SC
G
S
B
skin surface
Stratum corneum; SC
Granular layer; G
Spinous layer; S
Basal layer; B
n
CARD14 mutation
(Inflammation)
skin surface
SC
S
B
m
Affected skin
Revertant spot #2
Revertant spot #3
n
Affected skin
Revertant spot #1
Revertant spot #2
Revertant spot #3
o
Normal skin
CARD14 mutation
CAPE skin
clonal expansion
Revertant spot
Chromosome 17
mutation
recombination
(LOH)
revertant cell

**Fig. S1. Clinical and histological features of Case 1 and 2.**

**a**-**c**, Generalised scaly erythema (**a**) and palmoplantar keratoderma (**b**, **c**) were observed in Case 1. Joint contractures of the fingers were also noted (**b**). **d**-**f**, Case 2 exhibited skin symptoms similar to Case 1. **g**-**k**, Both patients showed multiple normal-appearing spots. Skin biopsy was taken from the left forearm (**g**), left anterior chest (**h**), and right neck (**i**) of Case 1 and from the left thigh (**j**) and left lower leg (**k**) of Case 2. **l,** Normal skin structure and structural changes due to *CARD14* mutations. The epidermis (the outermost layer of the skin) is composed of layers of epithelial cells (keratinocytes). Beneath the epidermis lies the dermis comprising connective tissues and housing various appendages including blood vessels, hair follicles, sweat glands, and other structures. The normal epidermis has four layers: basal layer (B), spinous layer (S), granular layer (G), and stratum corneum (SC). Proliferative epidermal progenitors reside in basal layer, and all the cells above this layer are differentiated cells. Histology of the skin with *CARD14* mutations shows epidermal thickening (acanthosis), lack of a granular layer, and thickening of stratum corneum (hyperkeratosis) with retained nuclei (parakeratosis). **m, n**, Histological comparison between affected skin and revertant spots of Case 1. Haematoxylin and eosin stain. Scale bars, 100 μm. **o**, Schematic comparison among normal skin, CAPE skin, and revertant spot. The birth of a revertant cell via mitotic recombination and its clonal expansion lead to the appearance of clinically visible revertant spot.

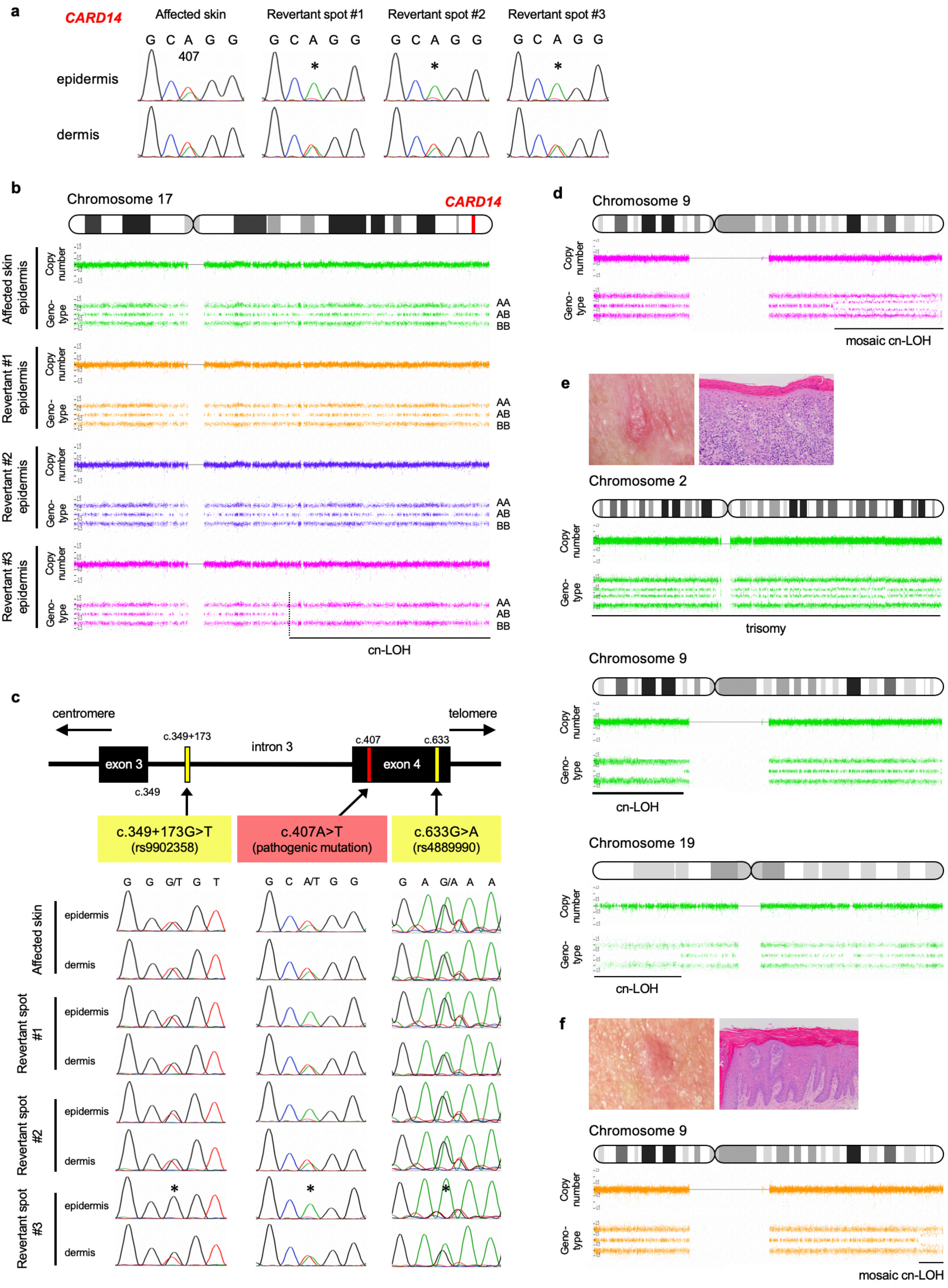
a
CARD14
Affected skin
Revertant spot #1
Revertant spot #2
Revertant spot #3
G C A G G
407
epidermis
dermis
b
Chromosome 17
CARD14
Affected skin epidermis
Revertant #1 epidermis
Revertant #2 epidermis
Revertant #3 epidermis
Copy number
Geno-type
AA
AB
BB
cn-LOH
c
centromere
telomere
c.349+173
intron 3
c.407
c.633
exon 3
exon 4
c.349
c.349+173G>T
(rs9902358)
c.407A>T
(pathogenic mutation)
c.633G>A
(rs4889990)
G G G/T G T
G C A/T G G
G A G/A A A
Affected skin
Revertant spot #1
Revertant spot #2
Revertant spot #3
d
Chromosome 9
mosaic cn-LOH
e
Chromosome 2
trisomy
Chromosome 9
cn-LOH
Chromosome 19
cn-LOH
f
Chromosome 9
mosaic cn-LOH

**Fig. S2. Genetic features of Case 1 and 2.**

**a**, Mutation analysis of *CARD14* using genomic DNA extracted from skin samples in Case 2. The heterozygous missense mutation, c.407A>T, is absent in the revertant epidermis (*). **b**, SNP array data of chromosome 17 from Case 2. LOH was identified in 1 of 3 revertant spots. Dotted lines represent the breakpoint of recombination. **c**, Heterozygous SNPs (c.349+173G>T and c.633G>A) on both sides closest to the pathogenic mutation in Case 2 were absent in the epidermis of revertant spot #3 (*) but were still present in the epidermis of revertant spots #1 and #2. Back mutation most likely explains the genetic reversion in revertant spots #1 and #2. **d**, Revertant spot #2 of Case 1 also harbours mosaic cn-LOH on chromosome 9. **e**, **f**, Clinical, pathological, and genetic features of SCC in situ arising on the left neck (**e**) and the posterior neck (**f**) of Case 1.

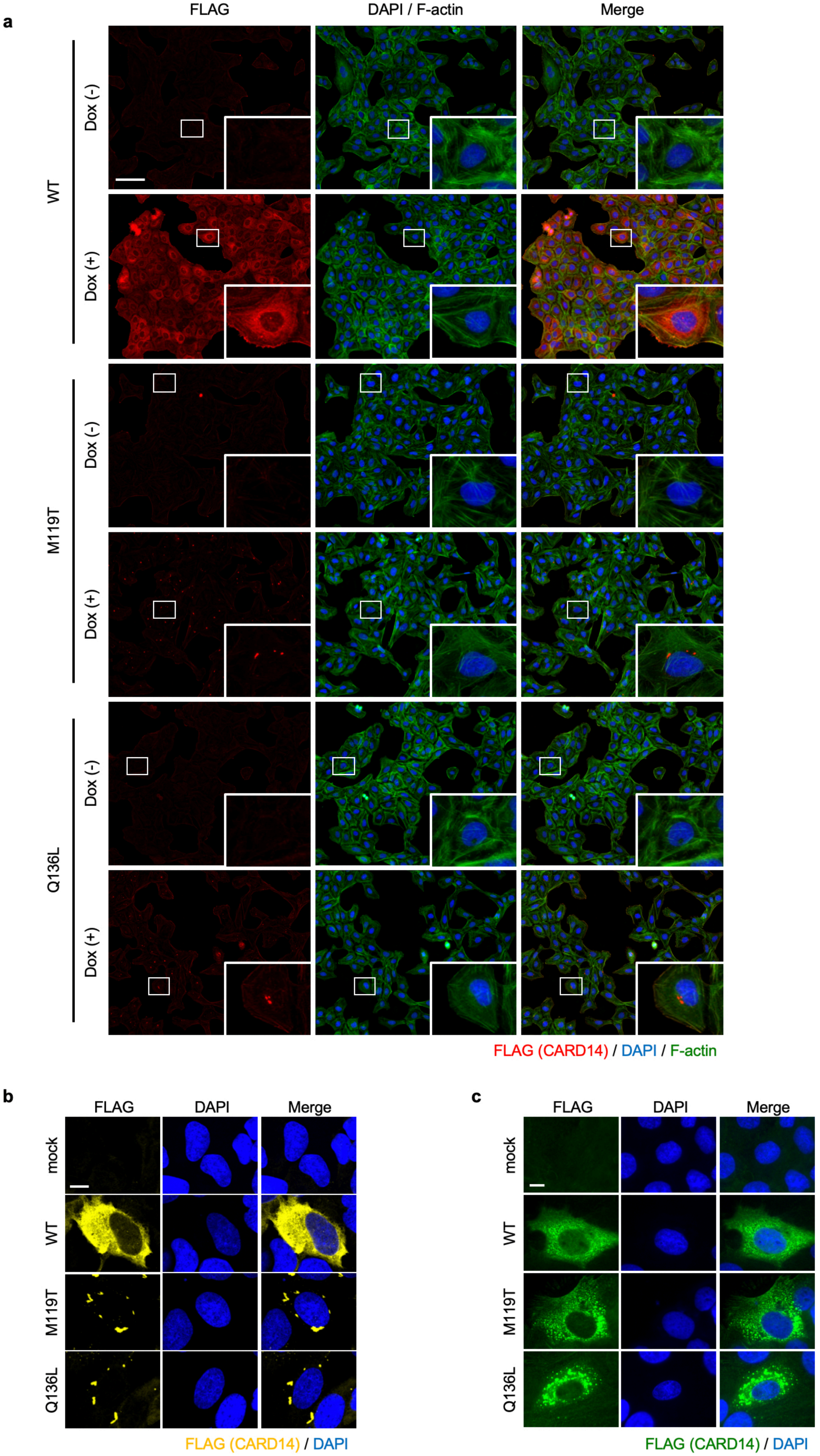
a
FLAG
DAPI / F-actin
Merge
WT
Dox (-)
Dox (+)
M119T
Dox (-)
Dox (+)
Q136L
Dox (-)
Dox (+)
FLAG (CARD14) / DAPI / F-actin
b
FLAG
DAPI
Merge
mock
WT
M119T
Q136L
FLAG (CARD14) / DAPI
c
FLAG
DAPI
Merge
mock
WT
M119T
Q136L
FLAG (CARD14) / DAPI

**Fig. S3. Immunofluorescence microscopy revealing intracellular CARD14 localisation.**

**a**, Immunofluorescence of CARD14 cell lines incubated with or without Dox for 24 h. Insets represent a high magnification of the area indicated in the boxes. Scale bar, 100 μm. **b**, **c**, Aberrant distribution of mut-CARD14 was further confirmed by transient overexpression analyses using U2OS cells (**b**) and HaCaT cells (**c**). Scale bar, 10 μm.

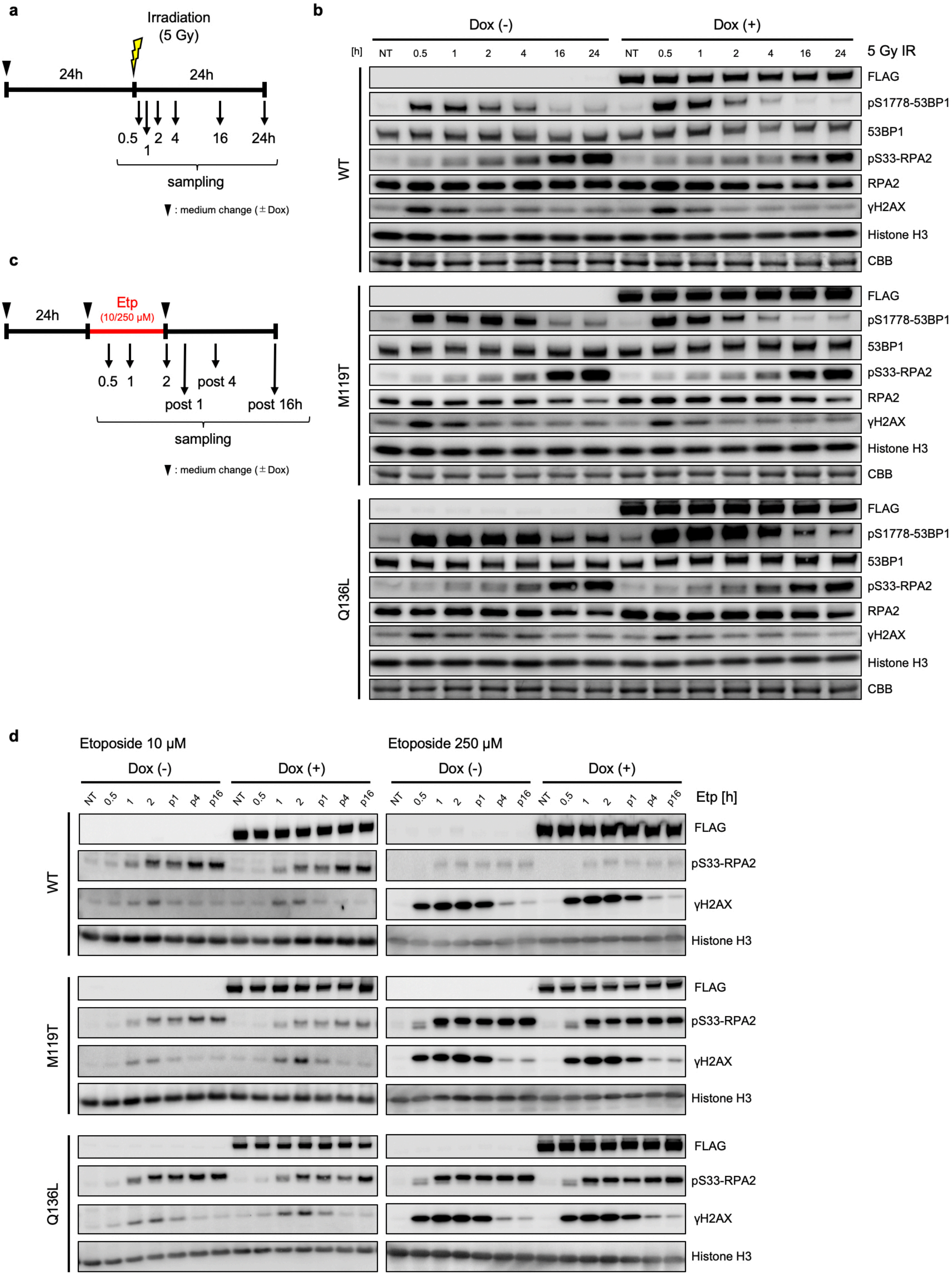

a
Irradiation (5 Gy)
24h
24h
0.5 1 2 4 16 24h
sampling
▼ : medium change (± Dox)
b
Dox (-)
Dox (+)
[h] NT 0.5 1 2 4 16 24 NT 0.5 1 2 4 16 24
5 Gy IR
WT
M119T
Q136L
FLAG
pS1778-53BP1
53BP1
pS33-RPA2
RPA2
γH2AX
Histone H3
CBB
c
24h
Etp (10/250 μM)
0.5 1 2 post 1 post 4 post 16h
sampling
▼ : medium change (± Dox)
d
Etoposide 10 μM
Etoposide 250 μM
Dox (-)
Dox (+)
NT 0.5 1 2 p1 p4 p16
Etp [h]
FLAG
pS33-RPA2
γH2AX
Histone H3

**Fig. S4. Mut-CARD14 expression provides no difference in the DDR.**

**a**, Schematic depicting timeline of when cells were exposed to 5 Gy IR. Whole-cell lysate was prepared at indicated times. **b**, Immunoblots showing the levels of DNA damage and the activation of 53BP1 and RPA2 following irradiation. pS1778-53BP1, phospho-Ser1778 53BP1. pS33-RPA2, phospho-Ser33 RPA2. NT, non-treatment. **c**, Schematic depicting timeline of when cells were exposed to 10 or 250 μM etoposide (Etp) and when the samples were obtained. **d**, Whole-cell lysate was immunoblotted with the indicated antibodies. p1/4/16, post 1/4/16 h. NT, non-treatment.

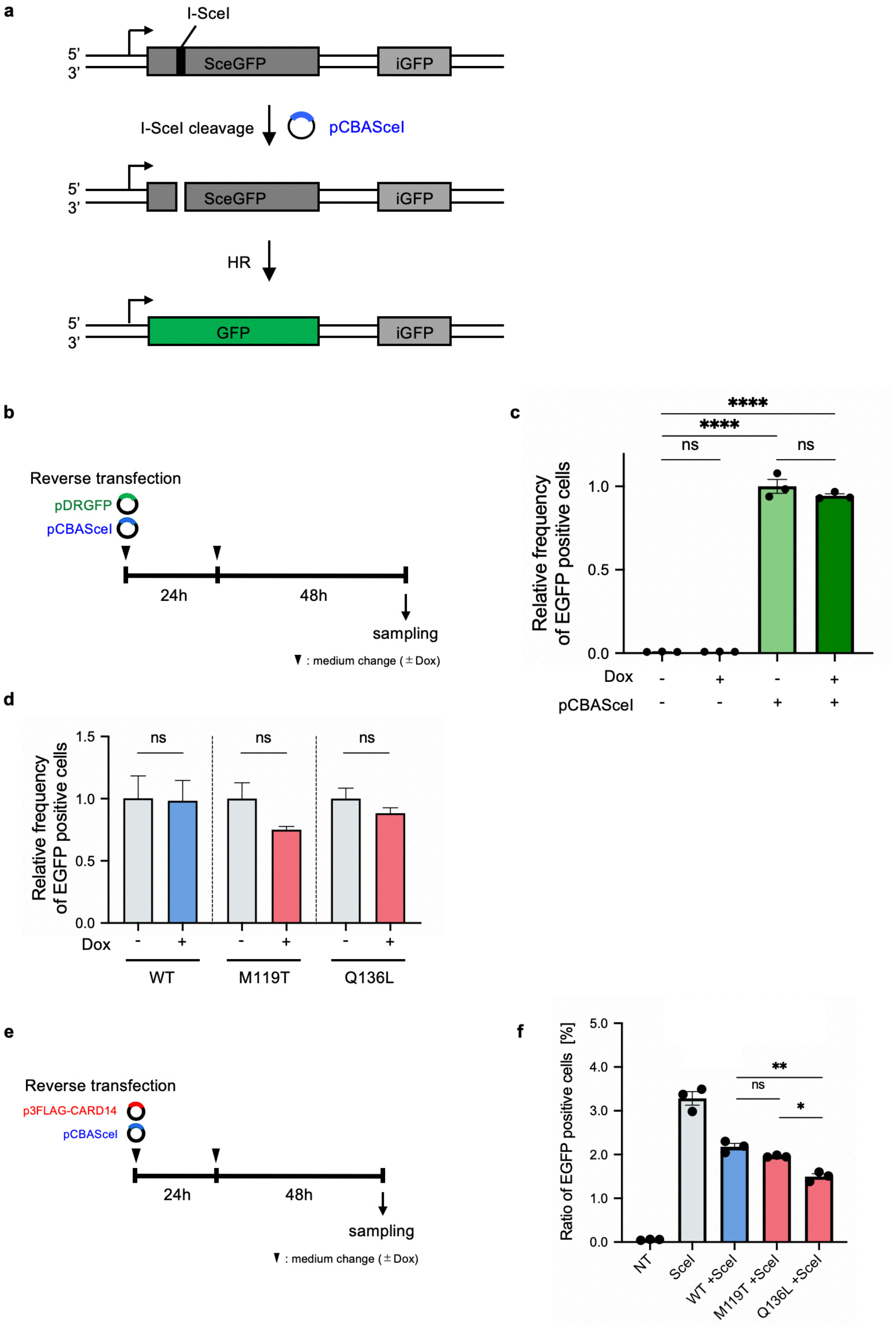

a
I-SceI
5'
3'
SceGFP
iGFP
I-SceI cleavage
pCBASceI
HR
GFP
b
Reverse transfection
pDRGFP
pCBASceI
24h
48h
sampling
▼ : medium change (± Dox)
c
****
****
ns
ns
Relative frequency of EGFP positive cells
1.0
0.5
0.0
Dox - + - +
pCBASceI - - + +
d
ns
ns
ns
1.5
1.0
0.5
0.0
Relative frequency of EGFP positive cells
Dox - + - + - +
WT
M119T
Q136L
e
Reverse transfection
p3FLAG-CARD14
pCBASceI
24h
48h
sampling
▼ : medium change (± Dox)
f
Ratio of EGFP positive cells [%]
5.0
4.0
3.0
2.0
1.0
0.0
**
ns
*
NT
SceI
WT +SceI
M119T +SceI
Q136L +SceI

**Fig. S5. Measurement of HR-mediated DSB repair frequency.**

**a**, Schematic showing the process of EGFP expression in DR-GFP assay. *SceGFP* gene contains I-SceI endonuclease site and cellular I-SceI expression leads to a DSB at this site. The DSB can be repaired by HR using downstream wild-type GFP sequence (iGFP) as a template, resulting in EGFP expression. **b**, **c**, Schematic depicting timeline of DR-GFP assay and when transient transfection of the DR-GFP reporter (pDRGFP) and I-SceI (pCBASceI) vectors was used (**b**). EGFP positive cells did not appear when U2OS cells were transfected with only pDRGFP but appeared when transfected with both pDRGFP and pCBASceI (**c**). DR-GFP assay works well even in transient transfection system; n = 3 independent experiments; error bars show s.e.m. **d**, DR-GFP assay by transient transfection revealed that neither wt-CARD14 nor mut-CARD14 increased HR frequency; n = 3 independent experiments. Error bars show s.e.m. **e**, **f**, Schematic of DR-GFP assay using U2OS cells stably expressing the DR-GFP reporter (**e**). In this assay, pCBASceI ± p3FLAG-CARD14 were transiently transfected (**e**). Again, neither wt-CARD14 nor mut-CARD14 promoted HR (**f**); n = 3 independent experiments; error bars show s.e.m. NT, non-treatment. Statistical significance was calculated using one-way ANOVA followed by a multiple comparisons test. *$P < 0.05$, **$P < 0.01$, ****$P < 0.0001$; ns, not significant.

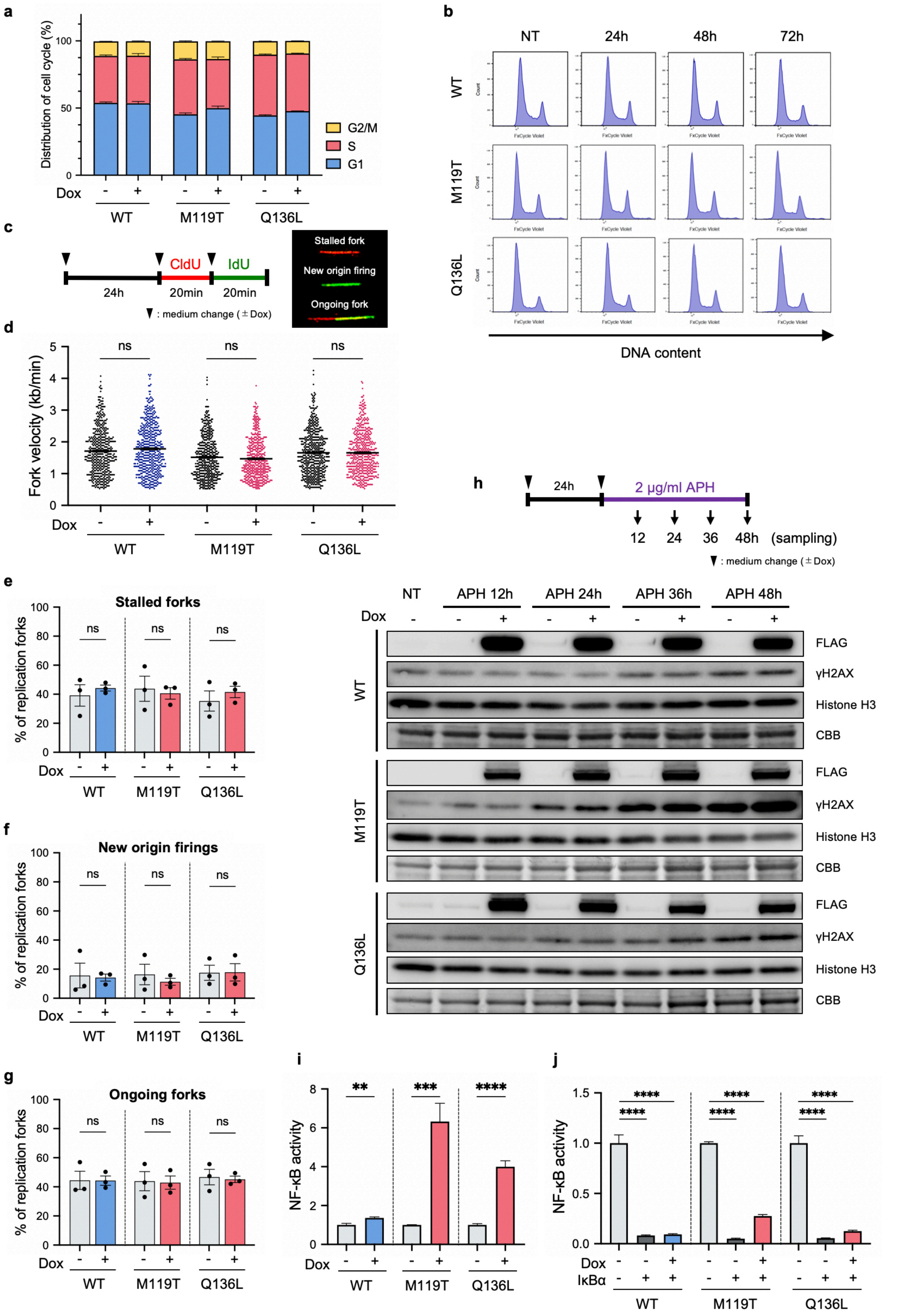

a
Distribution of cell cycle (%)
G2/M
S
G1
Dox
WT
M119T
Q136L
b
NT
24h
48h
72h
WT
M119T
Q136L
DNA content
c
CldU
IdU
24h
20min
20min
▼ : medium change (± Dox)
Stalled fork
New origin firing
Ongoing fork
d
Fork velocity (kb/min)
ns
Dox
WT
M119T
Q136L
e
Stalled forks
% of replication forks
ns
f
New origin firings
g
Ongoing forks
h
24h
2 μg/ml APH
12
24
36
48h
(sampling)
▼ : medium change (± Dox)
NT
APH 12h
APH 24h
APH 36h
APH 48h
FLAG
γH2AX
Histone H3
CBB
i
NF-κB activity
**
***
****
j
IκBα

**Fig. S6. Effect of mut-CARD14 expression on DNA replication.**

**a**, Cell cycle profiles of CARD14 cell lines at 24 h following Dox addition. For each sample, 30,000 cells were analysed; n = 3 independent experiments; error bars show s.e.m. **b**, Representative histograms of DNA content of CARD14 cell lines 24, 48, 72 h following Dox addition. **c**, Schematic depicting timeline of how cells are labelled with CldU and IdU. Representative images of a stalled fork, new origin firing, and ongoing fork are also shown. **d**, Comparison of replication fork velocity with or without Dox. At least 500 fibres for each cell line were quantified. Experiments were repeated at least thrice, and similar results were obtained each time. Horizontal lines represent mean values ± s.e.m. Statistical significance was calculated using the Mann-Whitney U test. ns, not significant. **e**, **f**, **g**, Quantification of the percentage of stalled forks (**e**), new origin firings (**f**), and ongoing forks (**g**) to all CldU-labelled forks; n = 3 independent experiments; error bars represent s.e.m. Statistical significance was calculated using the two-tailed t-test. **h**, Schematic depicting timeline of when cells were treated with 2 μg/ml APH and when samples were obtained. Whole-cell lysate was immunoblotted with indicated antibodies. Prolonged APH treatment increased γH2AX in the presence of mut-CARD14; NT, non-treatment. **i**, NF-κB-dependent luciferase assay using CARD14 cell lines incubated with or without Dox; n = 6 independent experiments; error bars show s.e.m. **j**, NF-κB-dependent luciferase assay using CARD14 cell lines to check the effect of IκBα overexpression; n = 6 independent experiments; error bars show s.e.m. Statistical significance was calculated using ordinary one-way ANOVA with multiple comparisons test. **$P < 0.01$, ***$P < 0.001$, ****$P < 0.0001$; ns, not significant.

a

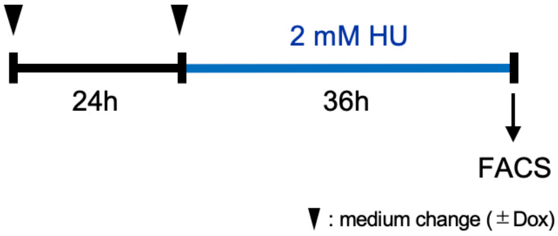
2 mM HU
24h
36h
FACS
: medium change (±Dox)

b

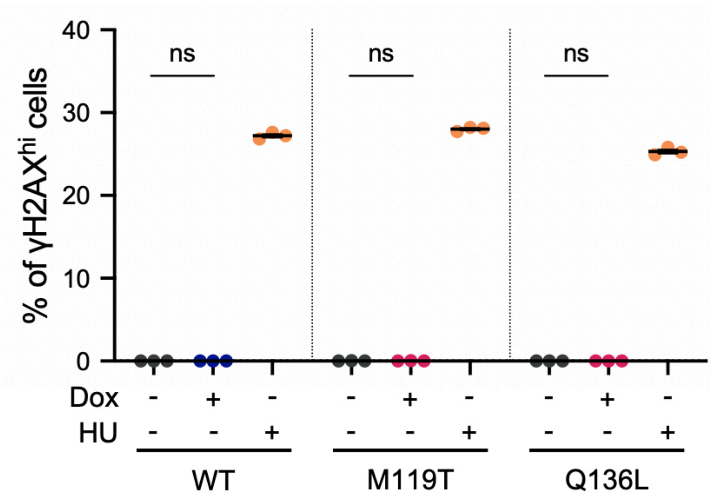
% of γH2AXhi cells
40
30
20
10
0
ns
ns
ns
Dox - + - - + - - + -
HU - - + - - + - - +
WT
M119T
Q136L

c

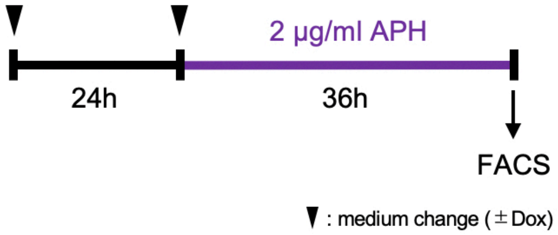
2 μg/ml APH
24h
36h
FACS
: medium change (±Dox)

d

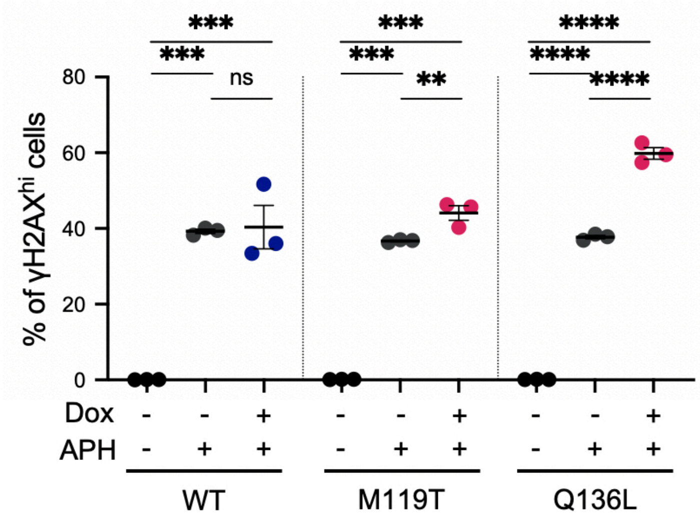
% of γH2AXhi cells
80
60
40
20
0
***
***
ns
***
***
**
****
****
****
Dox - - + - - + - - +
APH - + + - + + - + +
WT
M119T
Q136L

**Fig. S7. FACS-based analyses for evaluating the replication stress response.**

**a**, Schematic depicting timeline of when cells treated with HU for FACS analysis. **b**, CARD14 cell lines were incubated in the presence or absence of Dox for 36 h and harvested to estimate the levels of γH2AX by FACS. CARD14 expression per se did not increase the percentage of γH2AX$^{hi}$ cells. **c**, Schematic depicting timeline of when cells were treated with APH for FACS analysis. **d**, A clear increase in the percentage of γH2AX$^{hi}$ cells following APH treatment was observed in mut-CARD14 expressing cells. Statistical significance was calculated using ordinary one-way ANOVA with multiple comparisons test. **$P$ < 0.01, ***$P$ < 0.001, ****$P$ < 0.0001; ns, not significant.

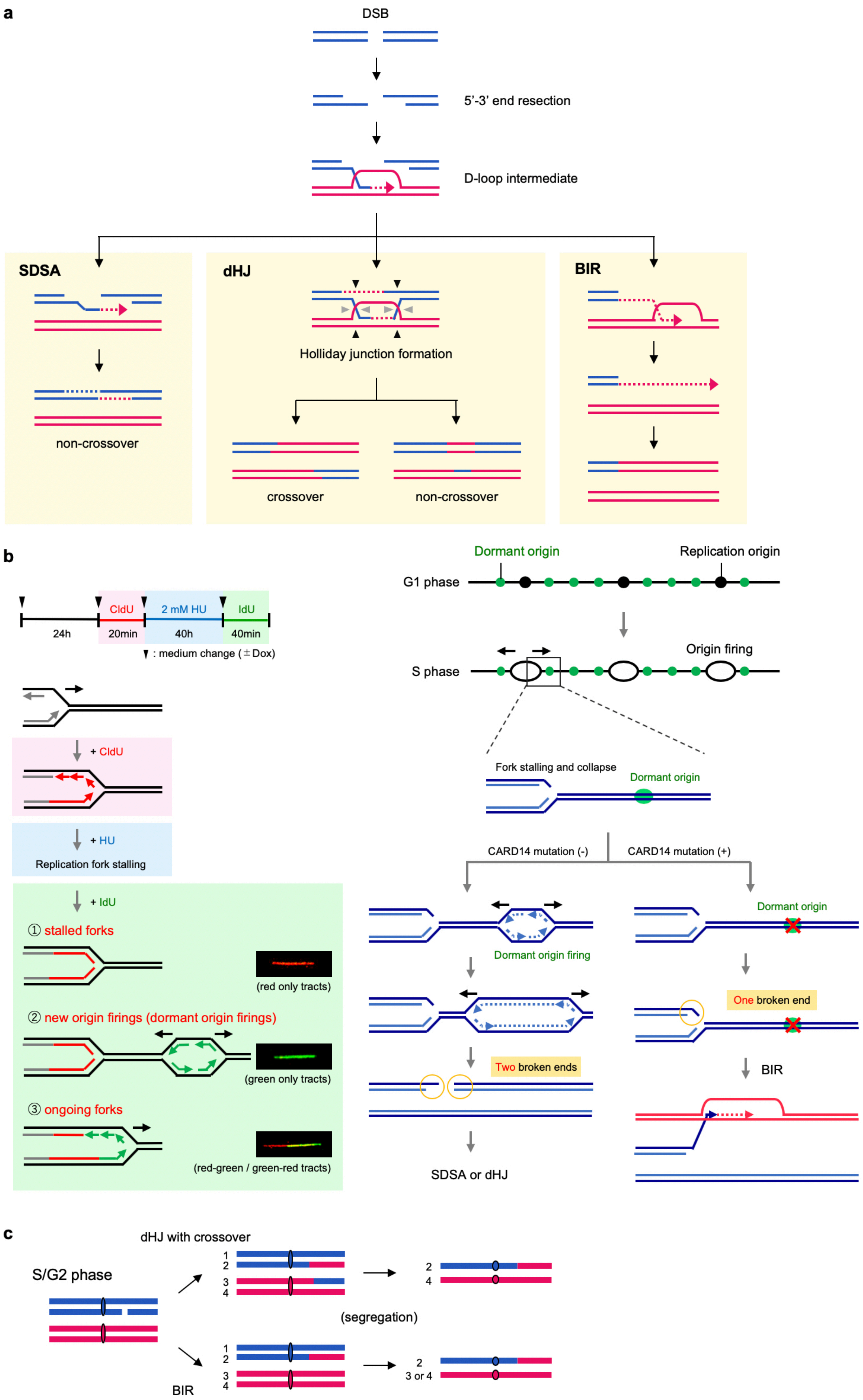
a
DSB
5'-3' end resection
D-loop intermediate
SDSA
non-crossover
dHJ
Holliday junction formation
crossover
non-crossover
BIR
b
CldU
2 mM HU
IdU
24h
20min
40h
40min
: medium change (±Dox)
+ CldU
+ HU
Replication fork stalling
+ IdU
① stalled forks
(red only tracts)
② new origin firings (dormant origin firings)
(green only tracts)
③ ongoing forks
(red-green / green-red tracts)
Dormant origin
Replication origin
G1 phase
S phase
Origin firing
Fork stalling and collapse
Dormant origin
CARD14 mutation (-)
CARD14 mutation (+)
Dormant origin firing
Dormant origin
Two broken ends
One broken end
BIR
SDSA or dHJ
c
S/G2 phase
dHJ with crossover
1
2
3
4
2
4
(segregation)
BIR
1
2
3
4
2
3 or 4

**Fig. S8. Three different HR pathways repairing collapsed forks.**

**a**, HR initiates with a 5′-3′ end resection which produces a 3′ single-stranded end that invades a homologous template to form a displacement loop (D-loop) intermediate. All three sub-pathways can ensue from this D-loop structure and their outcomes are different. In SDSA, the nascent strand is displaced to anneal to the other 3′ single-stranded end, resulting in a non-crossover outcome. In the dHJ pathway, also referred to as DSB repair, the second DNA end is captured by the D-loop and a dHJ intermediate is formed. The following resolution can generate either a crossover (cleavage at black arrowheads on one side and grey arrowheads) or a non-crossover (cleavage at black or grey arrowheads) outcome. In BIR, extensive DNA synthesis from the invading end occurs when only one broken end is available for repair. **b**, Schematic showing the treatment of the cells, and representative images of three types of fibres are shown on the left. Suppression of dormant origin activation can cause the situation in which only one broken end is available for repair. **c**, Long-tract LOH observed in RM can be mediated by two HR pathways, BIR or dHJ, with a crossover outcome. LOH of entire distal regions of chromosomes may arise when DSBs are repaired by these two pathways in S/G2 phase and recombinant sister chromatids segregate to different daughter cells in the M phase.

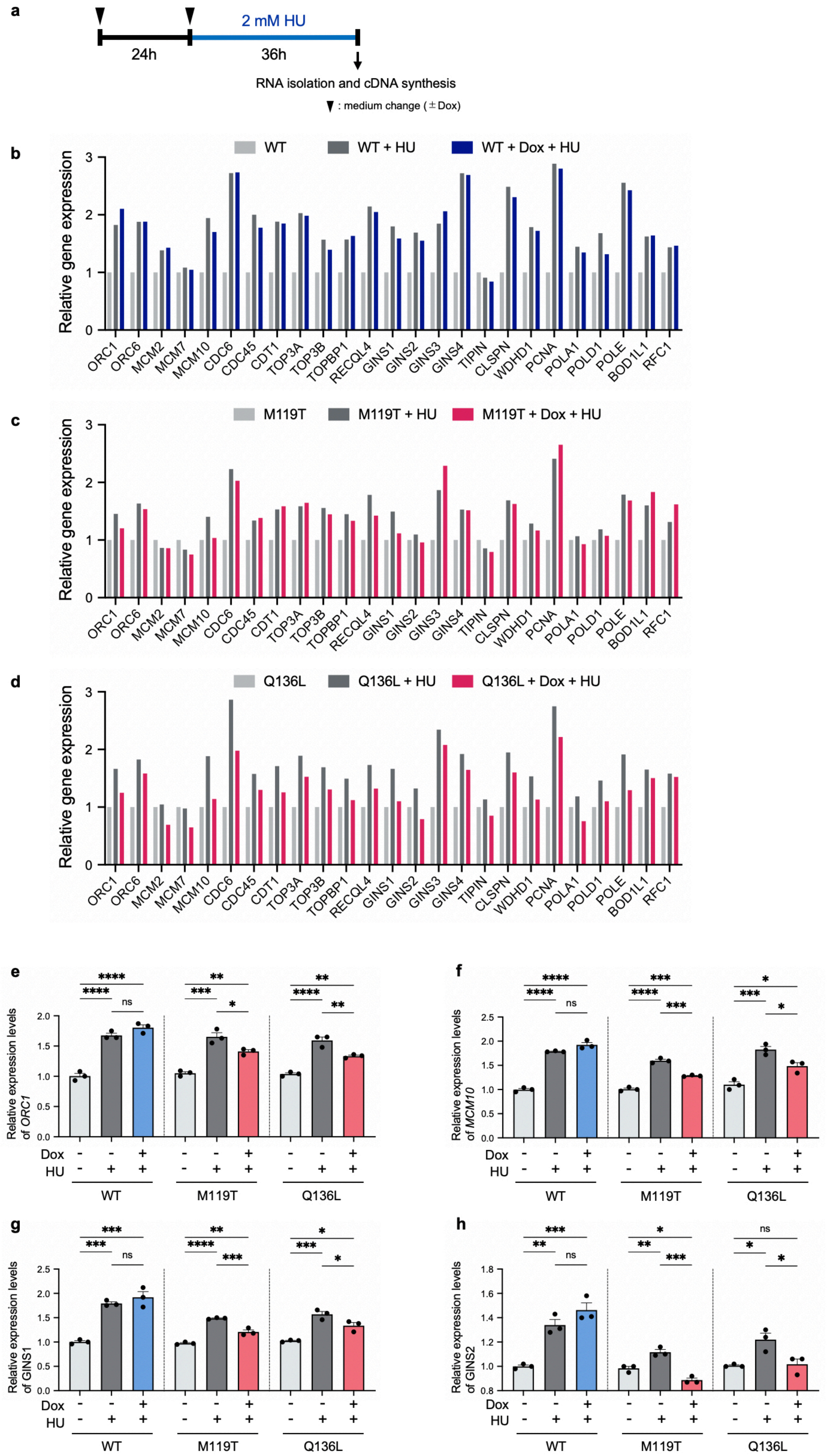
a
2 mM HU
24h
36h
RNA isolation and cDNA synthesis
▼ : medium change (± Dox)
b
WT
WT + HU
WT + Dox + HU
Relative gene expression
ORC1 ORC6 MCM2 MCM7 MCM10 CDC6 CDC45 CDT1 TOP3A TOP3B TOPBP1 RECQL4 GINS1 GINS2 GINS3 GINS4 TIPIN CLSPN WDHD1 PCNA POLA1 POLD1 POLE BOD1L1 RFC1
c
M119T
M119T + HU
M119T + Dox + HU
Relative gene expression
d
Q136L
Q136L + HU
Q136L + Dox + HU
Relative gene expression
e
Relative expression levels of ORC1
f
Relative expression levels of MCM10
g
Relative expression levels of GINS1
h
Relative expression levels of GINS2
Dox
HU
WT
M119T
Q136L
ns

**Fig. S9. qPCR screening analysis of replication-related genes revealed the suppressive state of new origin firing in mutant CARD14 expressing cells.** **a-d**, RNA was isolated from CARD14 cell lines which were incubated in the presence or absence of Dox for 24 h and with or without 2 mM HU for 36 h (**a**). Gene expression levels were analysed by qPCR (n =1 biological replicate, each with 2 technical replicates) (**b-d**). Note that expression values were normalised to ACTB expression. **e-h**, Gene expression levels of *ORC1* (**e**), *MCM10* (**f**), *GINS1* (**g**), and *GINS2* (**h**) were further analysed using the newly prepared samples; n = 3 independent experiments; error bars show s.e.m; Statistical significance was calculated using ordinary one-way ANOVA with multiple comparisons test. *$P < 0.05$, **$P < 0.01$, ***$P < 0.001$, ****$P < 0.0001$; ns, not significant.
